# Disease tolerant mice shelter a pathogenic intestinal microbiota able to trigger lethal disease in low tolerant hosts

**DOI:** 10.1101/2025.02.20.639353

**Authors:** Vinícius M. Vidal, Letícia M. Ferreira, Bahtiyar Yilmaz, Elena Montes-Cobos, Andreza M.S. Gama, Jamil Z. Kitoko, Samara G. Rosa, Gabriel A. Teixeira, José N. D. Farias, Varsha Raghavan, Dan Littman, Heitor S.P. de Souza, Fábio B. Canto, Marcelo T. Bozza

## Abstract

Disease tolerance is a defensive strategy that limits tissue damage during infection. Macrophage migration inhibitory factor (MIF)-deficient mice (*Mif^-/-^*) are protected in different models of infection and intestinal inflammation due to unclear disease tolerance mechanisms, whereas low disease-tolerant *Il10^-/-^*mice develop microbiota-dependent spontaneous gut inflammation. Here, we examined whether IL-10 is required for the phenotype seen in *Mif^-/-^*mice and, conversely, the contribution of MIF during IL-10 deficiency. While breeding for double-deficient *Mif^-/-^Il10^-/-^* mice, we unexpectedly observed that *Il10^-/-^* individuals died within days after co-housing with *Mif^-/-^* mice. We found that healthy *Mif^-/-^*hosts endure a highly diverse, unique and dysbiotic-like microbiota composition, including antibiotic-resistant *Enterobacteriaceae* species, which were sufficient to cause acute and lethal Th1-driven colitis in *Il10^-/-^* recipients. The disease was characterized by increased frequencies of IFNγ^+^ cells and neutrophils within colonic lamina propria. *Mif^-/-^Il10^-/-^*mice died prematurely, and survivors developed communicable disease, - indicating that lack of IL-10 is a dominant trait. These findings suggest that tolerant individuals harbor a gut microbiota enriched in pathogens/pathobionts, which can trigger disease in susceptible hosts.

## Introduction

Host-microbe interactions influence several biological processes in metazoans, shaping immune, metabolic and neural physiology, and ultimately regulating the host’s health status (Levy et al., 2017). The intricate network of interactions between host-symbiont and microbe-microbe, whether cooperative or competitive, is further modulated by environmental factors such as diet and exposure to xenobiotics (Culp et al., 2024; Koppel et al., 2017; Link et al., 2024). The composition of the microbiome at mucosal barriers is finely tuned by the host through immune-regulated mechanisms – *e.g.* mucus secretion, antimicrobial peptide production, CD4^+^Foxp3^+^ T cell activity and IgA/IgM production. Impairment or loss of these mechanisms disrupts the host’s ability to maintain health-related commensal colonization, thus favoring infection and inflammation (Kawamoto et al., 2014; Spragge et al., 2023; Vaishnava et al., 2011; Wang et al., 2015; Wlodarska et al., 2014).

Defense strategies against infection can be broadly classified as avoidance, resistance or disease tolerance (Medzhitov et al., 2012). Resistance mechanisms control infectious load, while disease tolerance mechanisms limit the damage caused either directly by the infectious agent, or indirectly by the immune response against it. Hence, disease tolerance preserves tissue and organic function regardless of pathogen burden (Medzhitov et al., 2012; Schneider and Ayres, 2008). Variations in disease tolerance among individuals with similar microbial load account for the differing outcomes observed during infection – ranging from asymptomatic cases and mild disease to severe illness or death.

Macrophage migration inhibitory factor (MIF) is a pleiotropic protein highly conserved across metazoan evolution. In mammals, MIF is recognized for its pro-inflammatory properties, promoting the production of other inflammatory mediators (Bozza et al., 1999; Calandra et al., 1994, 1995; Donnelly et al., 1997; Mitchell et al., 2002). MIF also retains macrophages and neutrophils in inflamed tissue (Bernhagen et al., 2007; Galvao et al., 2016; Takahashi et al., 2009), and provides survival signals to macrophages, T cells and B cells (Bacher et al., 1996; Gore et al., 2008; Hudson et al., 1999; Mitchell et al., 2002). Under steady-state conditions, MIF is abundant in the gastrointestinal (GI) tract mucosa, produced by cells in the epithelial monolayer, and immune cells in the lamina propria (Maaser et al., 2002; De Souza et al., 2015).

Humans with inflammatory bowel disease (IBD) exhibit elevated MIF plasma concentrations compared to both healthy individuals and patients receiving Infliximab treatment (De Jong et al., 2001). Furthermore, *Mif^-/-^* mice are protected from disease in various IBD models, indicating a pathogenic role of MIF in intestinal inflammation. *Mif^-/-^* mice are protected from dextran sulfate sodium (DSS)-induced colitis, showing increased survival rates and reduced numbers of macrophages and neutrophils in the affected tissue (Ohkawara et al., 2008). In the CD4^+^CD45RB^high^ T cell transfer model of colitis, double-deficient *Rag^-/-^Mif^-/-^* or *Rag^-/-^* hosts treated with α-MIF antibodies exhibit reduced body weight loss, decreased intestinal inflammation and lower amounts of inflammatory cytokines in the colon (De Jong et al., 2001). We have previously shown that *Mif^-/-^* mice are protected from oral *Toxoplasma gondii* infection compared to WT controls. These mice have improved survival and decreased intestinal inflammation, although displaying increased parasite load in the ileum (Cavalcanti et al., 2011). Similarly, α-MIF-treated WT mice survive *Clostridium difficile* infection despite the sustained bacterial burden (Jose et al., 2018). These results indicate that MIF contributes to tissue damage during gut inflammation and its absence enhances disease tolerance mechanisms, promoting survival.

IL-10, a key regulatory protein in mucosal tissues, is essential for maintaining intestinal homeostasis and preventing excessive inflammation. Humans that carry loss-of-function mutations in the IL-10 receptor gene (*IL10R*) experience very early-onset inflammatory bowel disease (VEO-IBD) (Ashton et al., 2017; Dong et al., 2021; Sandy et al., 2022), while *Il10^-/-^* mice similarly develop spontaneous intestinal inflammation (Kühn et al., 1993). Disease in *Il10^-/-^* mice is dependent on the presence of commensal micro-organisms, as germ-free mice do not develop pathology (Sellon et al., 1998). Conventionally-raised *Il10^-/-^* mice also show earlier disease onset compared to specific pathogen-free (SPF) mice (Kühn et al., 1993). This indicates that exposure to pathobionts/pathogens triggers intestinal inflammation in the absence of IL-10 signaling. In fact, *Il10^-/-^* mice succumb to enteropathogenic microorganisms such as murine norovirus (MNV) (Basic et al., 2014), *Helicobacter hepaticus* (Kullberg et al., 1998, 2001, 2006)*, Citrobacter rodentium* (Krause et al., 2015) and oral infection by *T. gondii* (Gazzinelli et al., 1996). In these models, *Il10^-/-^* mice display heightened gut inflammation and tissue damage, despite having similar or even reduced infectious loads compared to WT mice. These results underscore the protective effect of IL-10 in gut inflammation and highlight the low disease-tolerance status of individuals lacking this regulatory cytokine.

Given that high-tolerant *Mif^-/-^* and low-tolerant *Il10^-/-^* mice exhibit opposing phenotypes in the gut mucosa, we questioned which one would dominate in a double-deficient *Mif^-/-^Il10^-/-^* mouse. Experiments with *Mif^-/-^Il10^-/-^* mice and respective controls will help clarify the specific role of MIF on tissue damage and gut inflammation observed in conventionally raised *Il10^-/-^* mice. Simultaneously, these studies will also determine the contribution of IL-10 to the resilience of *Mif^-/-^* mice against gut inflammation. During animal breeding, we unexpectedly noticed that *Il10^-/-^*mice died few days after living with *Mif^-/-^* cohabitants. This observation led us to hypothesize that *Mif^-/-^* mice, due to their high tolerance to gut disease, might be harboring a microbiota enriched with pathobionts and pathogens able to trigger disease in *Il10^-/-^* individuals.

## Results

### *Il10^-/-^* mice develop early and severe colitis after co-housing with *Mif^-/-^* mice

To systematically investigate the initial observation that *Il10^-/-^* mice succumb when living with *Mif^-/-^* cohabitants, we set up different co-housing conditions, pairing *Il10^-/-^* individuals with either *Mif^-/-^* or WT mice (Figure 1A). *Il10^-/-^* mice began to progressively lose weight after six days of co-housing with *Mif^-/-^* mice and died within two weeks (Figure 1B). These animals exhibited typical signs of intestinal disease - *i.e.* rectal prolapse, and soft stool containing visible blood (Figure 1C). One week after co-housing onset, colonoscopy evidenced focal points of intestinal inflammation characterized by hyperemia, edema, hemorrhages and ulcerative lesions (Figure 1D). Macroscopically, the colons of diseased *Il10^-/-^* mice displayed redness, scarce presence of fecal pellets and shorter length (Figure 1E). Histopathological analyses of colonic tissue sections revealed focal areas of inflammation, marked by epithelial erosion, ulcerative lesions, crypt abscess, and extensive cellular infiltrate in both the mucosal and submucosal regions (Figure 1F).

**Figure 1.**
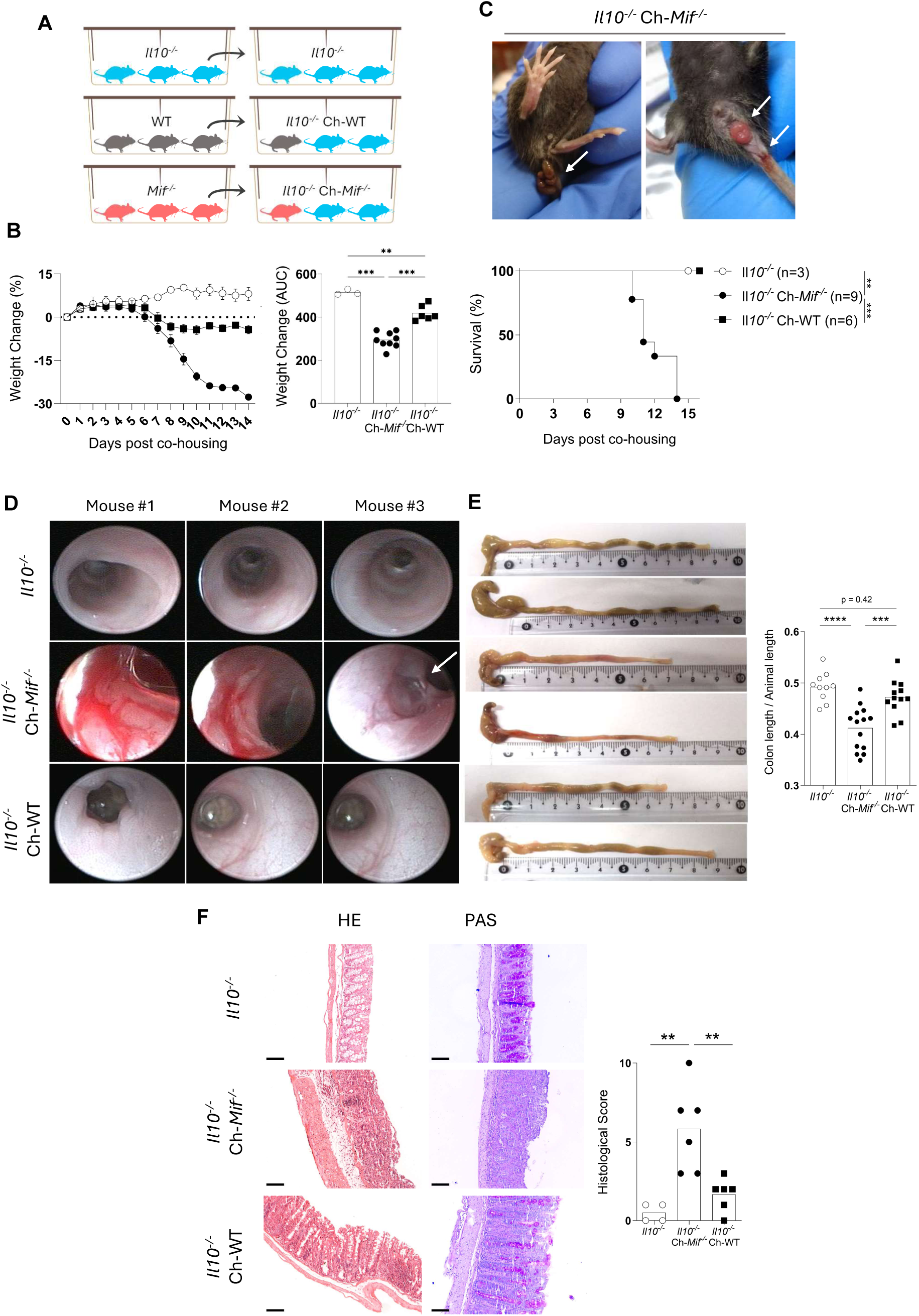
*Il10^-/-^* mice develop early and severe colitis after co-housing with *Mif^-/-^* mice. **(A)** Protocol scheme. *Il10^-/-^* mice were co-housed either with *Mif^-/-^*or WT mice in a 1:1 or 2:1 proportion. **(B)** *Il10^-/-^*mice body weight and survival curve within two weeks. Weight change was determined by the following equation: %n = ((Wn / W0) -1) * 100, where Wn = mouse weight on day N, and W0 = mouse weight on day 0. Mean ± SEM. Mice were euthanized if 25% of initial body weight was lost. **(C)** Clinical signs of intestinal inflammation in *Il10^-/-^* mice co-housed with *Mif^-/-^* mice. Blood in the stool and rectal prolapse are depicted in the arrows. **(D)** Colonoscopy images of *Il10^-/-^* mice after seven days of co-housing. Each picture belongs to a different mouse and is representative of at least five mice analyzed in each experimental group. White arrow: ulcer. **(E)** Macroscopic view of *Il10^-/-^* mice large intestine after ten days of co-housing. Photographs depict colon length, the presence of stool and tissue redness. **(F)** Colon photomicrographs of *Il10^-/-^* mice after ten days of co-housing. Hematoxylin/Eosin (HE), and Periodic Acid-Schiff (PAS) staining were performed on 5 µm colon sections. Scale Bar = 100 µm; magnification = x100. The histopathological score considers epithelial injury, infiltrating leukocytes, and the presence of ulcers and erosions. See Experimental Procedures for a description of the scoring algorithm. **(B)** Representative data from three independent experiments. **(E and F)** Pool of two independent experiments. One-way ANOVA, post-hoc Tukey. Survival curves were compared with log-rank Mantel-Cox test. * for adjusted p ≤ 0.05; ** for adjusted p ≤ 0.01; *** for p ≤ 0.001; **** for p ≤ 0.0001.

In contrast, *Il10^-/-^* mice co-housed with WT controls did not present clear signs of gut inflammation or develop lethal disease (Figure 1B-F). While initially failing to gain weight as non-co-housed *Il10^-/-^* counterparts, they eventually recovered and survived throughout the two-month observation period (Figure S1A). Moreover, we found no sex bias in this phenomenon, as both male and female *Il10^-/-^* mice developed colitis when co-housed with either male or female *Mif^-/-^* animals (Figure S1B). From the perspective of WT and *Mif^-/-^* mice, neither group experienced weight loss (Figure S1C) or showed any sign of disease while co-housed with *Il10^-/-^* mice. Moreover, the disease completely depended on IL-10 deficiency, since WT mice co-housed with *Mif^-/-^* individuals did not lose weight (Figure S1D).

To confirm that *Mif^-/-^* mice in our colony do not exhibit low-grade colon inflammation, we collected tissue samples from them under steady-state conditions. Similar to WT controls, *Mif^-/-^* mice displayed preserved colonic tissue architecture and no signs of inflammation (Figure S2A). The expression levels of the pro-inflammatory cytokines *Tnfa* and *Il6* were comparable between both genotypes (Figure S2B). Additionally, *Mif^-/-^* mice bred and gained weight normally in our facility (Figure S2C). These findings indicate that *Il10^-/-^* mice develop severe colitis and ultimately succumb when co-housed with *Mif^-/-^* individuals.

### *Il10^-/-^* mice exhibit an increase in IFNγ^+^CD4^+^ T cells following co-housing with *Mif^-/-^* mice

To identify the cells and mediators responsible for driving inflammation in the colons of *Il10^-/-^*mice co-housed with the *Mif^-/-^* genotype, we performed immunophenotyping of colonic lamina propria and epithelium, in addition to mesenteric lymph nodes, using flow cytometry (Figure S3). TNFα, a key inflammatory cytokine in the intestinal mucosa, is a target for immunotherapy in IBD patients (Noor et al., 2024). *Il10^-/-^* mice co-housed with either *Mif^-/-^* or WT groups showed an increase of TNFα^+^ cells in the colonic lamina propria (Figure S4A), suggesting that TNFα may not be the primary mediator distinguishing the effects of co-housing.

We also observed that *Il10^-/-^* mice co-housed with *Mif^-/-^* animals exhibited an increase in IFNγ-producing cells (Figure S4B), particularly IFNγ^+^CD4^+^ T lymphocytes in the colonic lamina propria (Figure 2A) and mesenteric lymph nodes (Figure 2B). Interestingly, both CD4^+^Foxp3^-^ effector Th1 cells and CD4^+^Foxp3^+^ Treg cells produced IFNγ in diseased *Il10^-/-^* mice (Figure 2A, B). Despite their increase in the colonic lamina propria (Figure S5A) and mesenteric lymph nodes (Figure S5B) of these mice, CD4^+^Foxp3^+^ Treg cells appear insufficient in number and/or function to control the rampant inflammation. In fact, in *Il10^-/-^* mice co-housed with *Mif^-/-^* mice the CD4^+^Foxp3^+^ Treg cells may become subverted, likely contributing to the inflammation by secreting TNFα, IFNγ and IL-17A (Figure 2A, B and S5).

**Figure 2.**
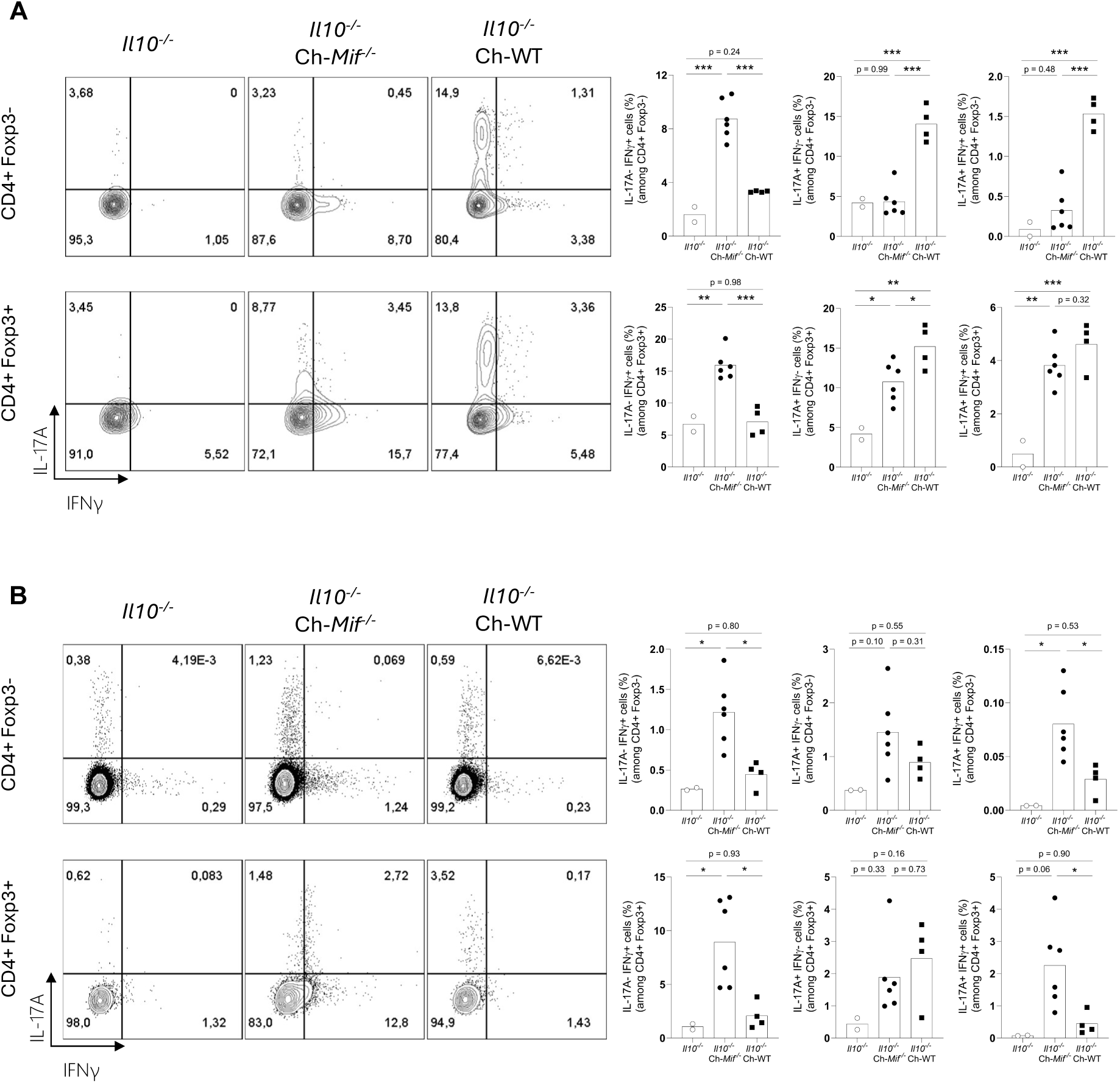
*Il10^-/-^* mice display increased IFNγ^+^CD4^+^ T cells after co-housing with *Mif^-/-^* mice. Flow cytometry analysis of *Il10^-/-^* mice after ten days of co-housing. **(A)** colonic lamina propria and **(B)** mesenteric lymph nodes. Relative frequencies of IFNγ^+^ and IL-17A^+^ cells among CD4^+^Foxp3^-^ and CD4^+^Foxp3^+^ cells. Cells were gated after doublet, debris and dead cell exclusion. See Figure S2 for gating strategies and FMO controls. Representative contour plots from each experimental group. One-way ANOVA, post-hoc Tukey. * for adjusted p ≤ 0.05; ** for adjusted p ≤ 0.01; *** for p ≤ 0.001.

Within the myeloid compartment, *Il10^-/-^* mice co-housed with *Mif^-/-^* counterparts showed an extensive neutrophil infiltrate (CD11b^+^Ly6G^+^SSC^high^) (Figure S6A) in the colon. In contrast, no significant differences were observed in eosinophil (CD11b^+^Ly6G^-^SiglecF^+^SSC^high^) frequencies (Figure S6B). However, there was a marked reduction of myeloid dendritic cells (CD11b^+^Ly6G^-^Ly6C^-^F4/80^-^CD11c^+^) (Figure S6C), monocytes (CD11b^+^Ly6G^-^Ly6C^+^F4/80^-^) and macrophages (CD11b^+^Ly6G^-^F4/80^+^) (Figure S6D).

The co-housing with WT controls elicited a Th17 skewed response in *Il10^-/-^* mice (Figure 2A, B and S4A), marked by an increase of IL-17A^+^CD4^+^ T lymphocytes in the colon, but not in the mesenteric lymph nodes (Figure 2A, B). These mice did not exhibit significant changes in the frequencies of CD4^+^Foxp3^+^ Treg cells (Figure S5) or myeloid cells (Figure S6). Taken together, these findings suggest that the disease induced in *Il10^-/-^* mice through co-housing with *Mif^-/-^* individuals is primarily driven by the pathogenic actions of IFNγ^+^CD4^+^ T cells and neutrophils.

### *Mif^-/-^* mice harbor a distinct fecal microbiota enriched in pathobionts/pathogens, that is sufficient to induce colitis in *Il10^-/-^* hosts

To test the hypothesis that sharing of fecal microbiota during co-housing triggered disease, we performed daily transfers of fecal pellets from the *Mif^-/-^* or WT donors to those of *Il10^-/-^* recipients. Only *Il10^-/-^* mice exposed to the fecal pellets derived from *Mif^-/-^* donors exhibited marked weight loss, succumbing over three weeks (Figure 3A). We also performed a single fecal transfer via oral gavage, and, once again, only the fecal suspension from *Mif^-/-^* mice was lethal to *Il10^-/-^* animals (Figure 3B). The filtered fecal suspension of *Mif^-/-^* donors did not induce disease in *Il10^-/-^* mice (Figure 3B), suggesting that pathogenesis was not driven by a virus or soluble factors, such as a toxin, but rather by microorganisms larger than 0.22 µm present in the feces.

**Figure 3.**
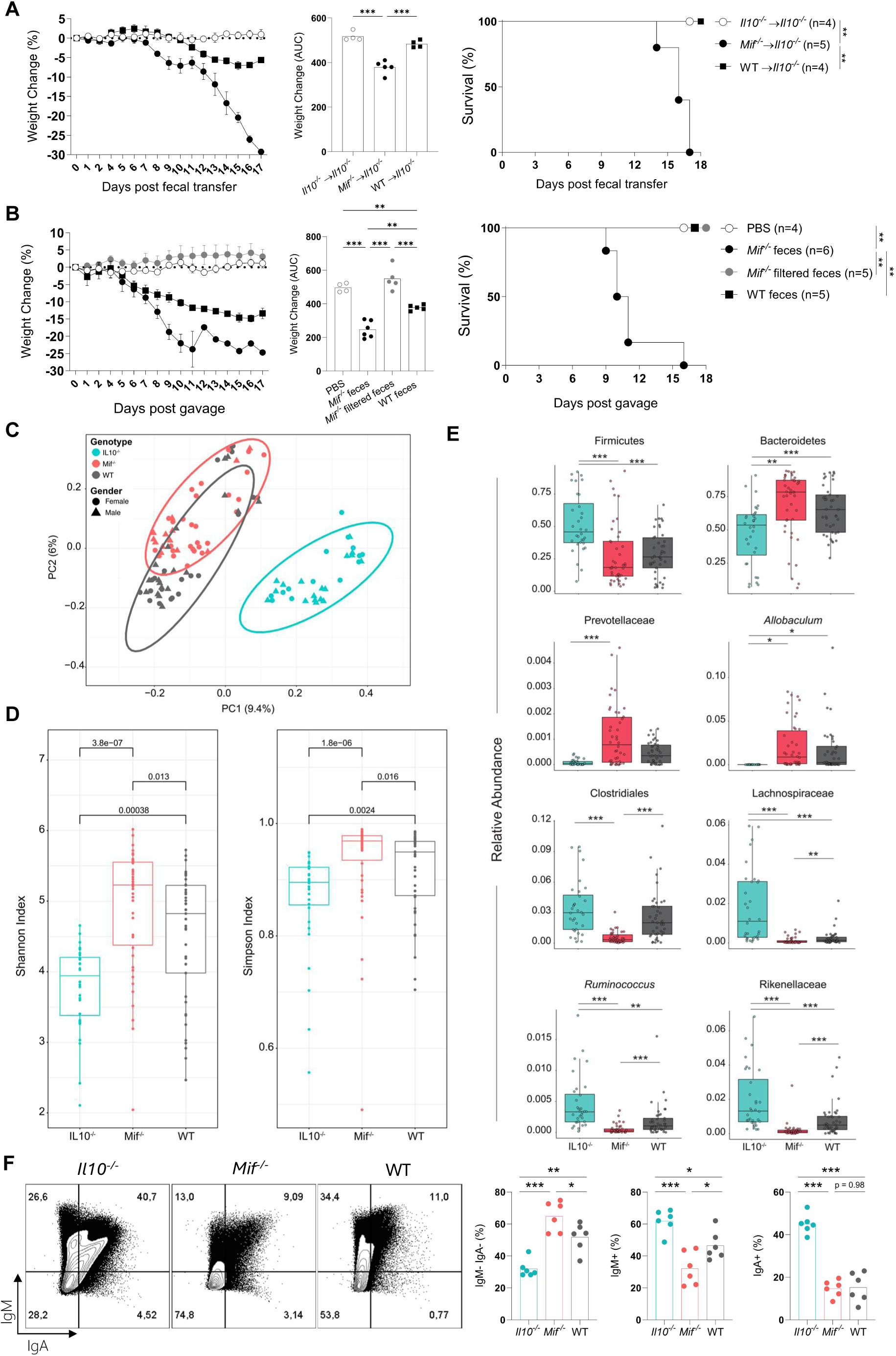
*Mif^-/-^* mice shelter a distinct and highly diverse fecal microbiota sufficient to trigger lethal disease in *Il10^-/-^* mice. **(A and B)** *Il10^-/-^* mice body weight (left) and survival (right) curves after fecal transfer. Mean ± SEM. **(A)** Fecal pellets from *Mif^-/-^* or WT donors were daily transferred to *Il10^-/-^* recipients cages. (B) *Il10^-/-^* mice received a single fecal transplant from *Mif^-/-^* or WT mice donors through oral gavage. Filtered samples were obtained using a 0.22 μm membrane filter. **(C-E)** Fecal samples from *Mif^-/-^*, *Il10^-/-^* and WT mice under steady state conditions were analyzed for microbiome composition based on the 16S rRNA gene sequencing. **(C)** Principal Coordinate Analysis (PCoA) plot of β-diversity using UniFrac distance matrix. **(D)** Species richness (α-diversity) using Shannon and Simpson indices. **(E)** Differential taxa abundance among genotypes. **(F)** Bacterial cells were isolated from fresh fecal pellets from *Mif^-/-^*, *Il10^-/-^* and WT mice under steady state conditions. Frequencies of IgA and IgM coated bacteria by flow cytometry. Representative contour plots. One-way ANOVA, post-hoc Tukey. Survival curves were compared with log-rank Mantel-Cox test. * for adjusted p ≤ 0.05; ** for adjusted p ≤ 0.01; *** for p ≤ 0.001.

We investigated the microbiota composition in fresh fecal pellets collected from WT, *Mif^-/-^* and *Il10^-/-^* mice under steady-state conditions using 16S rRNA gene sequencing. The analysis of β-diversity revealed that each genotype harbored a unique microbiota profile (Figure 3C). Although there was partial clustering overlap between *Mif^-/-^* and WT mice, their microbiomes were significantly distinct. Regarding α-diversity, *Mif^-/-^* mice had the highest species diversity, while *Il10^-/-^* animals showed the lowest (Figure 3D). In the gut, high α-diversity is typically associated with a healthy microbiome (Clooney et al., 2021; Franzosa et al., 2018); however, *Mif^-/-^* mice harbored what appeared to be a dysbiotic microbiota despite their increased diversity (Figure 3E). The microbiota colonizing *Mif^-/-^* mice exhibited the lowest Firmicutes/Bacteroidetes ratio (Figure 3E and S7A) and was enriched in bacteria such as *Prevotellaceae* and *Allobaculum* spp. (Figure 3E and S7B, C), which have been linked to intestinal inflammation. In contrast, health-associated taxa, including Clostridiales, *Lachnospiracea*, *Rikenellaceae,* and *Ruminococcus* spp., were more abundant in the *Il10^-/-^*group, but were scarcely represented in *Mif^-/-^* mice (Figure 3E and S7B, C). In this context, while the microbiota colonizing *Il10^-/-^* mice exhibited low species diversity, it contained a high abundance of beneficial commensal bacteria typically associated with a healthy status. These findings suggest that *Mif^-/-^* mice harbor a microbiota enriched in pathobionts/pathogens sufficient to trigger lethal disease in *Il10^-/-^* animals. The fact that *Mif^-/-^* individuals can maintain a healthy phenotype despite this potentially pathogenic microbiota implies that MIF impairs tolerance to disease in the colonic mucosa.

### *Mif^-/-^* mice harbor a community of bacteria largely uncoated by immunoglobulins

Host-derived factors, including antibodies, actively shape microbiota composition. Previous studies have identified high IgA coating as a hallmark of colitogenic bacteria in IBD (Palm et al., 2014). Thus, we analyzed IgA and IgM coating of bacteria in fresh fecal samples, harvested from WT, *Mif^-/-^* and *Il10^-/-^* donors, using flow cytometry (Figure S8A). *Mif^-/-^* mice exhibited the highest frequency of double-negative IgM^-^IgA^-^ (uncoated) bacteria, primarily due to a reduction of IgM^+^ cells, while *Il10^-/-^* animals harbor a community of bacteria highly coated by IgM and IgA (Figure 3F and Figure S8B). This suggests that MIF might influence microbiota composition through distinct patterns of antibody production.

### Antibiotic-resistant *Klebsiella oxytoca* and *Proteus mirabilis* strains harvested from *Mif^-/-^* donors elicit lethal disease in *Il10^-/-^* recipients

Considering the possibility that disease-causing bacteria colonizing the gut of *Mif^-/-^* mice were responsible for the lethal outcome in IL-10 deficient animals, we treated *Mif^-/-^* and/or *Il10^-/-^* mice daily via oral gavage with the antibiotic cocktail AVNM (Ampicilin, Vancomycin, Neomycin and Metronidazole) for ten days prior to the co-housing experiment, and maintained the treatment throughout the study. However, AVNM treatment did not prevent disease transmission to *Il10^-/-^* mice, which still presented weight loss and clear signs of colitis, eventually succumbing after co-housing with *Mif^-/-^* animals (Figure 4A, B).

**Figure 4.**
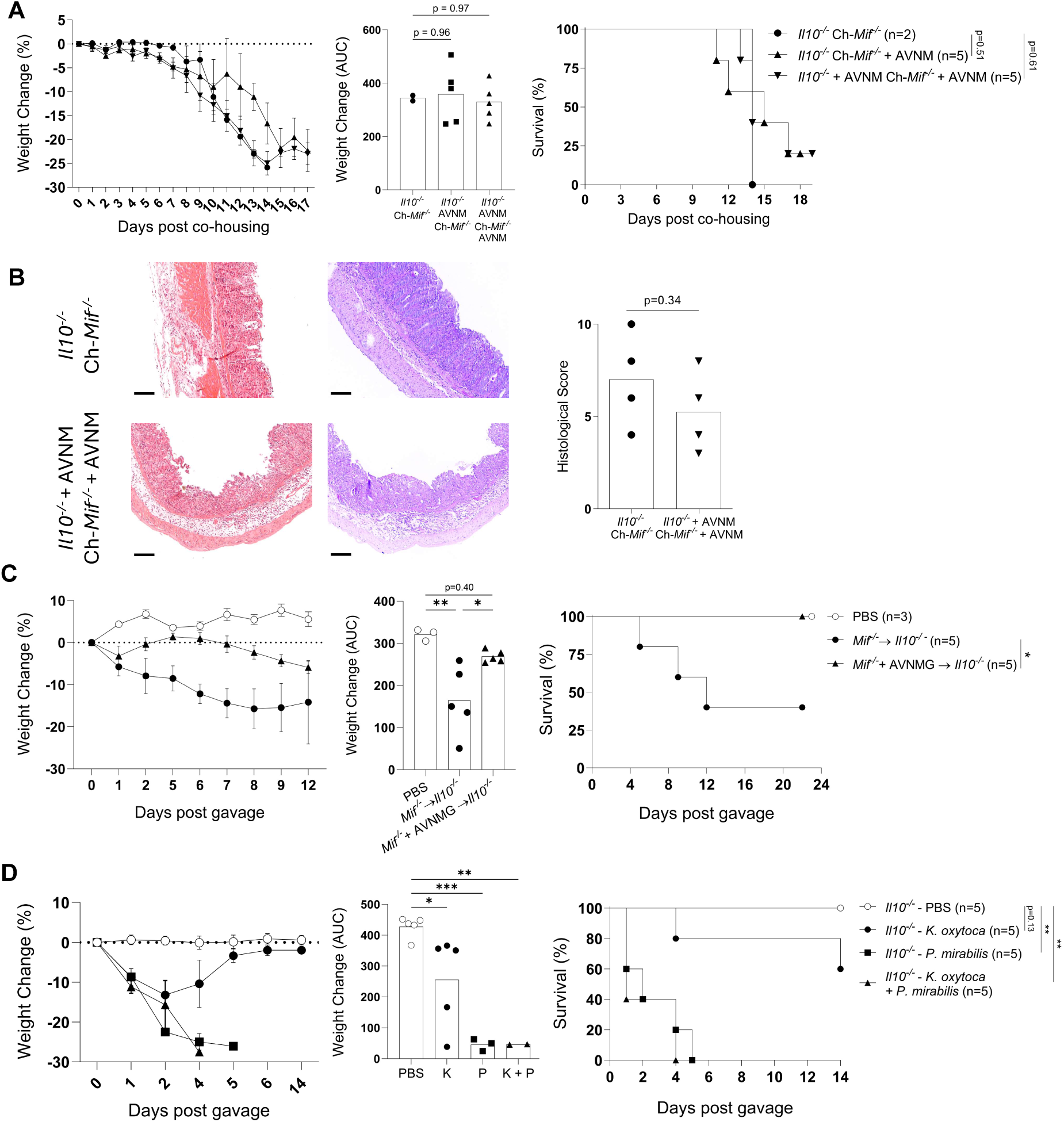
Antibiotic-resistant *Klebsiella oxytoca* and *Proteus mirabilis* strains harvested from *Mif^-/-^* donors elicit lethal disease in *Il10^-/-^* recipients. **(A and B)** *Il10^-/-^* and/or *Mif^-/-^* mice were pre-treated daily through gavage with antibiotic cocktail AVNM (Ampicillin, Neomycin, Metronidazole and Vancomycin) for ten days before co-housing. The treatment was maintained daily throughout the experiment. (A) *Il10^-/-^* mice body weight (left) and survival (right) curves after co-housing. Representative data from three independent experiments. **(B)** Hematoxylin/Eosin (HE), and Periodic Acid-Schiff (PAS) staining were performed on 5 µm colon sections. Scale Bar = 100 µm; magnification = x100. The histopathological score considers epithelial injury, infiltrating leukocytes and the presence of ulcers and erosions. Two-tailed T-test. See experimental procedures for scoring. (C) *Mif^-/-^* mice were treated with a gentamycin-containing antibiotic cocktail (AVNMG) for ten days in drinking water. Enriched bacterial fraction from fresh fecal samples was transferred to *Il10^-/-^* through oral gavage. *Il10^-/-^* mice body weight (left) and survival (right) curves after fecal transplant. (D) *Il10^-/-^* mice were mono-inoculated through oral gavage with 1 x 10^9^ CFU of *P. mirabilis*, 1 x 10^9^ CFU of *K. oxytoca,* or co-inoculated with 0.5 x 10^9^ CFU of each species. Mice body weight (left) and survival (right) curves after inoculum. **(A, C, D)** Weight mean ± SEM. One-way ANOVA, post-hoc Tukey. Survival curves were compared with log-rank Mantel-Cox test. * for adjusted p ≤ 0.05; ** for adjusted p ≤ 0.01; *** for p ≤ 0.001.

To test whether the disease-causing agent was a bacterium resistant to the AVNM cocktail, we collected fecal samples from *Il10^-/-^* mice after one week of co-housing with *Mif^-/-^* counterparts (both under AVNM treatment), cultured the samples under anaerobe conditions, and identified the isolates through partial 16S rRNA gene sequencing. Indeed, we isolated *Klebsiella oxytoca*, *Proteus mirabilis*, *Enterococcus* spp. and *E. coli*/*Salmonella* (Figure S9A), which were resistant to the AVNM but susceptible *in vitro* to Gentamycin (Figure S9B). We then treated *Mif^-/-^* mice with an antibiotic cocktail that included Gentamycin (AVNMG) for ten days, collected fecal samples, and transferred a bacteria-enriched fraction via oral gavage to *Il10^-/-^*recipients. AVNMG treatment in *Mif^-/-^* mice inhibited the lethal disease observed in *Il10^-/-^* hosts following fecal transfer (Figure 4C).

Antibiotic-resistant *P. mirabilis* and *K. oxytoca* have been linked to intestinal inflammation in both humans and experimental models (Garrett et al., 2010; Högenauer et al., 2006; Zhang et al., 2021). Indeed, *Il10^-/-^* mice that were either mono-inoculated with *P. mirabilis* or *K. oxytoca,* or co-inoculated intragastrically with both species developed acute lethal disease (Figure 4D). In contrast, the inoculum with these species was less lethal to WT hosts (Figure S10). Altogether, our data suggest that *Mif^-/-^* mice harbor antibiotic-resistant *P. mirabilis* and *K. oxytoca* capable of triggering lethal disease in *Il10^-/-^* hosts.

### Double-deficient *Mif^-/-^Il10^-/-^* mice die prematurely and survivors develop communicable disease

To test whether MIF blockade would have a protective effect on intestinal inflammation associated with IL-10 deficiency, male *Il10^-/-^* and female *Mif^-/-^* mice were initially crossed to establish a double-deficient colony. From the breeding of F1 mice, which were heterozygous for both loci and phenotypically healthy, we selected F2 *Mif^-/-^Il10^+/-^*animals as parents to continue crossings. Out of 245 mice in the F3 offspring, we only observed 37 *Mif^-/-^Il10^-/-^* mice (15,1%), significantly deviating from the expected distribution (χ² [2, 0.05] = 17.376, two-tailed p = 0.0002) (Figure 5A). Most of the double-deficient mice exhibited reduced life expectancy (median = 9.4 weeks, Q1 = 4.5, Q3 = 17.2), did not gain weight as expected and developed visible signs of disease, such as rectal prolapse (Figure 5B). Histopathological analysis of their colons revealed extensive inflammation, characterized by intense cellular infiltrate, epithelial erosion, and crypt abscess formation (Figure 5B).

**Figure 5.**
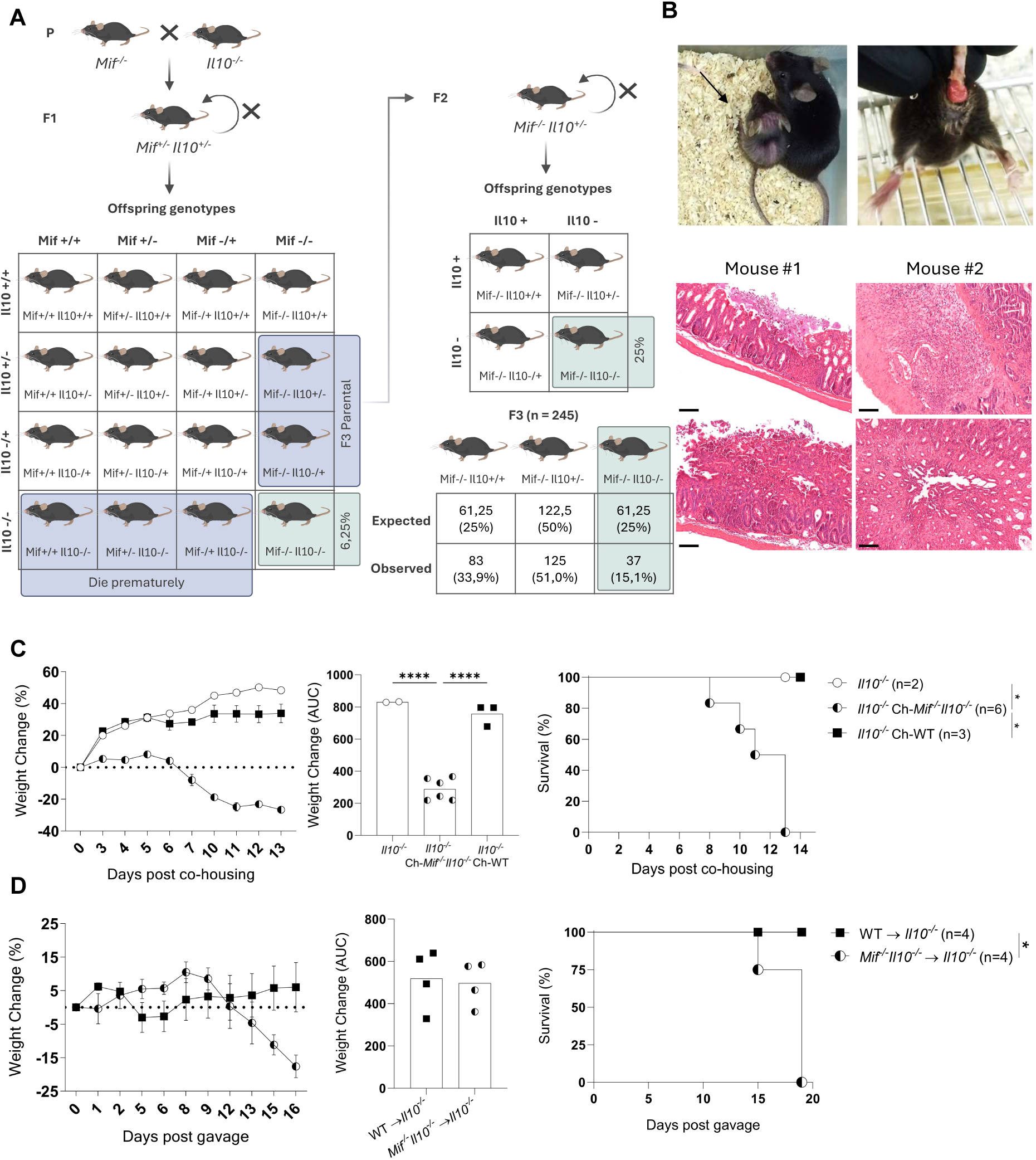
Double-deficient *Mif^-/-^Il10^-/-^* mice die prematurely, and survivors develop communicable disease. **(A)** Punnet square diagram and breeding strategy for obtaining double-deficient *Mif^-/-^Il10^-/-^* mice. **(B)** Representative photographs of a *Mif^-/-^Il10^-/-^* mouse (black arrow) and a littermate displaying signs of disease and intestinal inflammation. Hematoxylin/Eosin (HE) staining was performed on 5-µm colon sections. Scale Bar = 100 µm; magnification = x100. **(C)** *Il10^-/-^* mice body weight (left) and survival (right) curves after co-housing *Il10^-/-^Mif^-/-^* mice in a 3:1 proportion respectively. Weight mean ± SEM. One-way ANOVA, post-hoc Tukey. **(D)** *Il10^-/-^* mice received a single fecal transplant through oral gavage from *Il10^-/-^Mif^-/-^* or WT mice donors. Weight mean ± SEM. Two-tailed T-test. **(C,D)** Survival curves were compared with log-rank Mantel-Cox test. * for adjusted p ≤ 0.05; **** for p ≤ 0.0001.

*Il10^-/-^* mice co-housed with *Mif^-/-^Il10^-/-^* animals experienced progressive weight loss and died within two weeks (Figure 5C). Fecal transfer from *Mif^-/-^Il10^-/-^*donors via oral gavage also induced lethal disease in *Il10^-/-^* recipients (Figure 5D). Additionally, *Il10^-/-^* mice were treated daily with neutralizing anti-MIF (α-MIF) antibodies before being challenged with co-housing. While the α-MIF treatment delayed disease onset, it failed to protect *Il10^-/-^* mice from lethal disease when co-housed with *Mif^-/-^* animals (Figure S11). Thus, blocking MIF signaling, either genetically or through antibody neutralization, was insufficient to prevent disease in *Il10^-/-^* mice.

## Discussion

Here, we identify MIF as a key regulator of the colonic microbial community, an inhibitory factor for disease tolerance and a promoter of tissue damage in the intestinal mucosa. *Mif^-/-^* mice harbor a potentially pathogenic microbiota capable of inducing wasting disease in genetically susceptible *Il10^-/-^* individuals. Contrary to expectations, impairment of MIF biological activity did not attenuate microbiota-driven inflammation in *Il10^-/-^* mice. This highlights the dominant role of IL-10 in maintaining mucosal disease tolerance and intestinal homeostasis.

Horizontal transfer of colitogenic bacteria via co-housing has been documented, typically involving ailing animals transferring disease to healthy counterparts. For example, double-deficient *Tbet^-/-^Rag2^-/-^*(TRUC) mice develop spontaneous colitis and transfer colitogenic microbiota that causes disease in WT controls (Garrett et al., 2007). Genetic deficiencies, such as those in *Nlrp6*, *Il18* and *Itch,* can alter microbiota composition and increase susceptibility to gut inflammation transferable to WT mice (Elinav et al., 2011; Kathania et al., 2020). We report a unique case of lethal colitogenic co-housing involving otherwise healthy *Mif^-/-^*mice.

*Il10^-/-^* mice developed different immune responses in the colonic mucosa after living with either WT or *Mif^-/-^* counterparts. *Il10^-/-^* mice co-housed with WT controls exhibited increased TNFα^+^ leukocytes and CD4^+^IL-17A^+^ T cells in the colon, but this was not sufficient to cause wasting disease. In contrast, *Il10^-/-^* individuals co-housed with *Mif^-/-^* mice displayed not only an increase in TNFα^+^ cells and a mild rise in CD4^+^IL-17A^+^ T cells, but also a unique expansion of CD4^+^ IFNγ^+^ T lymphocytes. Indeed, both IFNγ and IL-17 act synergistically to induce maximal inflammation in *Il10^-/-^* mice infected with *H. hepaticus* (Kullberg et al., 2006). We found that TNFα, IL-17A and IFNγ were significantly produced by Foxp3+ Treg cells, suggesting a loss of their suppressive function. IL-10, crucial for Foxp3 stability, is key to Treg function (Asseman et al., 1999; Chaudhry et al., 2011). Under inflammation, Tregs can lose Foxp3 expression (ex-Tregs), gaining the ability to produce IFNγ and IL-17A (Komatsu et al., 2014; Zhou et al., 2009). We also observed Foxp3 and pro-inflammatory cytokine coexpression in *Il10^-/-^* mice, similar to reports in *T. gondii*-infected WT mice (Oldenhove et al., 2009), suggesting a pathogenic shift in Tregs. Additionally, *Il10^-/-^* mice co-housed with *Mif^-/-^* animals exhibited a prominent neutrophil infiltrate, likely contributing pathologically to tissue damage. Neutrophil infiltration is a hallmark of active IBD, with its role potentially promoting or inhibiting inflammation depending on the context (Danne et al., 2023).

The microbiota of *Mif^-/-^* mice shares similarities with that of both IBD patients and experimental models associated with dysbiosis, notably an increased abundance of Bacteroidetes and reduction of Firmicutes (Man et al., 2011). These mice lack key short-chain fatty acid (SCFA) producers like *Lachnospiracea* and *Ruminococcus*, which are important for gut health (Arpaia et al., 2013; Furusawa et al., 2013; Parada Venegas et al., 2019). Clostridiales and *Rikenellaceae* were also underrepresented, while inflammatory-associated *Prevotellaceae* and *Allobaculum* spp. were abundant, resembling findings in *Itch^-/-^* and *Nlrp6^-/-^*mice. *Prevotella* spp. provokes metabolic changes in the microbiome, reducing acetate and IL-18 production, which increases susceptibility to DSS-induced inflammation (Iljazovic et al., 2021). *Allobacullum* spp. constitute highly IgA-coated colitogenic bacteria isolated from IBD patients (Palm et al., 2014) and *Allobacullum mucolyticum* was shown to degrade and feed on human mucins by secreting carbohydrate active enzymes (van Muijlwijk et al., 2021). Notably, *Mif^-/-^*mice tolerate *Enterobacteriaceae*, specifically *K. oxytoca* and *P. mirabilis*. *K. oxytoca* is a gram-negative pathobiont, found in 2-9% of healthy Europeans, and has been increasingly associated with intestinal inflammation (Beaugerie et al., 2003). *K. oxytoca*, linked to antibiotic-associated hemorrhagic colitis (AAHC), and *P. mirabilis*, known to disrupt epithelial integrity, (Hoffmann et al., 2010; Högenauer et al., 2006; Zhang et al., 2021) were sufficient to induce severe colitis in *Il10^-/-^* mice. *K. pneumoniae*, closely related to *K. oxytoca*, is linked to IBD in humans and mice, with strains from IBD patients inducing IFNγ-mediated colitis in germ-free *Il10^-/-^* mice (Federici et al., 2022). *K. pneumoniae* and *P. mirabilis* also cause colitis in TRUC mice and can transfer disease to WT mice (Garrett et al., 2010). Other isolates from *Mif^-/-^* mice, such as *Enterococcus spp.* and *E. coli*, could also worsen disease, as both have been linked to IBD in humans (Elhenawy et al., 2019; Seishima et al., 2019). Certain members of the microbiome, such as *Bacteroides* spp., *Helicobacter hepaticus* and *Bilophila wadsworthia*, are known to be particularly hazardous to *Il10^-/-^* mice (Bloom et al., 2011; Devkota et al., 2012; Kullberg et al., 2006; Xu et al., 2018). *Mif^-/-^* mice may harbor additional uncultured pathobionts.

SPF mice are valuable scientific tools but often fail to replicate human biology in part due to their limited environmental exposure and microbiota. Conventionally-raised mice, exposed to diverse microbial communities, better model human diseases and therapeutic interventions (Beura et al., 2016; Rosshart et al., 2019). In our facility, mice are frequently exposed to and colonized by parasites, like trichomonad protozoans and the nematode *Syphacia* spp., both of which can influence the bacterial community and modulate the host’s mucosal immune response. Notably, *Tritrichomonas* spp. colonization in WT mice induces CD4^+^IFNγ^+^ Th1 and CD4^+^IL-17A^+^ Th17 cells in the colon and aggravates T-cell-driven colitis in the *Rag1^-/-^*recipients (Chudnovskiy et al., 2016; Escalante et al., 2016; Gerrick et al., 2024). Conversely, *Syphacia* spp. has a protective effect during DSS-induced colitis by increasing Treg cells and IL-10 production (Taghipour et al., 2019). Although conventionally raised, *Il10^-/-^* mice in our facility thrive for up to one year without showing overt disease, indicating that their pathogen/pathobiont exposure is not extreme. This observation points to a unique interaction between the microbiota and host immune modulation, particularly influenced by MIF deficiency.

Although it is possible that double deficiency interfered with proper embryonic development, our findings suggest that *Mif^-/-^Il10^-/-^* mice likely died from environmental exposure to colitogenic microbiota from *Mif^-/-^* littermates. MIF and TNFα are cytokines that engage in mutual positive feedback loops (Calandra and Roger, 2003). It was shown that *Tnfr1^-/-^Il10^-/-^*double-deficient mice also develop spontaneous colitis early after weaning (Liu et al., 2020). In contrast to *Mif^-/-^* that show preserved tissue architecture with no overt signs of inflammation, nor increased expression of *Tnfa* and *Il6*, *Tnfr1^-/-^* animals exhibit intestinal abnormalities, including epithelial dysfunction in the colon, a higher frequency of neutrophils and IELs (Liu et al., 2020).

In humans, polarized intestinal epithelial cells (IECs) concentrate and release most of their intracellular MIF through the apical membrane (Maaser et al., 2002; de Souza et al., 2015). This release is rapidly triggered by infection with enteroinvasive *Salmonella dublin* or following exposure to IFNγ and IL-1α (Maaser et al., 2002). In the bobtail squid *Euprymna scolopes*, the MIF orthologue EsMIF is predominantly produced by epithelial cells in the invertebrate light organ. Its production is under circadian rhythm oscillation and orchestrates macrophage-like hemocyte migration into the light-organ crypts. Hemocytes periodically provide specific nutrients to the symbiont *Vibrio fischeri* that, in turn, adjust its metabolism accordingly to the presence/absence of hemocytes - indicating a critical and evolutionarily conserved role of MIF to engender robust host-microbial symbiosis (Koch et al., 2020).

Multicellular organisms have evolved alongside the ubiquitous, incessant and extremely diverse presence of microorganisms. Disease tolerance mechanisms allow harmonious interactions between hosts and symbionts (Ayres, 2016). Here, we provide experimental evidence that genetic deficiency in MIF enhances disease tolerance in the intestinal mucosa, allowing the host to harbor a microbiota enriched in pathobionts and/or pathogens while maintaining a healthy phenotype. While this microbiota is lethal to low tolerant individuals, such as those deficient in IL-10, it may confer benefits to the host. From an evolutionary perspective, disease tolerance increases the possibilities for host-symbiont interactions, thus expanding phenotypic plasticity and the chances of success under different environmental pressures. The pros and cons associated with maintaining a potentially pathogenic microbiota presumably depend on the context in which the host finds itself. Since humans display varying degrees of disease tolerance, our data might explain how apparently trivial social events, like cohabitation of high- and low-tolerant individuals sharing the same household, might set off inflammatory diseases such as IBDs.

## Methods

### Mice

Male and female C57BL/6 mice (*Mus musculus*), from six to twelve weeks-old, wild-type (WT), MIF-deficient (*Mif^-/-^*) previously described (Bozza et al., 1999) and backcrossed for 10 generations onto a C57BL/6 background, and IL-10-deficient (*Il10^-/-^*) (B6.129P2-Il10tm1Cgn/J -#002251 – Jackson Laboratories) were raised in conventional condition (non-SPF) in the Inflammation and Immunity Lab animal facility (IMPG / UFRJ). A health surveillance for circulating parasites, bacteria and viruses in the vivarium is provided in Supplementary Table 1. Mice were maintained in ventilated microisolator cages (Alesco, Monte Mor, Brazil), under controlled temperature conditions (22 ± 2 °C) and 12h light/dark cycle. The animals had free access to conventional irradiated food and sterilized water. All experimental procedures were approved by the Animal Ethics Committee (CEUA) (040/22 CEUA-UFRJ, Rio de Janeiro, Brazil).

### Co-housing

*Il10^-/-^* mice were co-housed either with WT, *Mif^-/-^* or *Mif^-/-^Il10^-/-^* mice in a 1:1, 2:1 or 3:1 proportion. Body weight change was determined by the following equation: %n = ((W_n_ / W_0_) -1) * 100, where W_n_ = body weight on day N, and W_0_ = body weight on day 0. Animals were euthanized if weight loss exceeded 25%.

### Fecal transfer

Roughly 20 fecal pellets were daily transferred from WT or *Mif^-/-^*donor cages to *Il10^-/-^* recipients cages. For the experiments using fecal transfer through oral gavage, 40 mg of feces were macerated in 1 mL of Phosphate Buffer Saline (PBS) and left for 1 min to decant. The supernatant was collected and diluted in 9 mL of PBS. Mice received 100 μL of the diluted fecal suspension in a single dose by gavage using the BD Angiocath plastic catheter holder (BD #388336). For bacterial enrichment, 90 mg of pooled feces from three mice were macerated in PBS and centrifuged at low speed (500 g, 5 min) and room temperature (RT). Supernatant was collected, washed and centrifuged again at low speed. Then, supernatant was collected and centrifuged at high speed (12000 g, 5 min, RT). The bacterial enriched pellet was washed twice and resuspended in 1 mL PBS. Inoculums of 100 μL of the suspension enriched in bacteria were administered by oral gavage.

### Histology

The colon was excised and cut transversely into three pieces. Feces were gently removed from the tube using curved forceps, and the pieces were fixed in buffered formaldehyde solution (4%) for 72h. Paraffin-embedded specimens were cut into 5-μm sections, and stained with Haematoxylin-Eosin (HE) and Periodic Acid-Schiff (PAS). Photomicrographs were taken using Olympus BX40 Microscope (Shinjuku, Tokyo, Japan) at ×100 magnification. The histopathological score was blindly performed and determined according to the following parameters: ulceration, hyperplasia, and inflammatory infiltrate. For ulceration, the scale considered was: 4, extensive and deep; 3, focal and deep; 2, extensive and superficial; 1, focal and superficial; 0, absent. For hyperplasia and inflammatory infiltrate, the scale considered was: 3, severe; 2, moderate; 1, light; 0, absent.

### Colonoscopy

After seven days of co-housing, mice were anesthetized in an isofluorane chamber and underwent colonoscopy with a bronchoscope (FB 120-P; Fujinon, Japan) coupled with a camera for recording.

### Antibiotic treatment

AVNM antibiotic cocktail was administered daily through oral gavage (100 μL) and contained anhydrous ampicillin (1 mg/mL) (Ampicilab), vancomycin hydrochloride (0.5 mg/mL) (ABL), neomycin sulfate (1 mg/mL) (Sigma #N6386), and metronidazole (1mg/mL) (Nidazofarma) diluted in PBS. AVNMG included gentamycin (1mg/mL) (Gentatec). Alternatively, antibiotic was provided in drinking water - ampicillin (0,08 mg/mL), vancomycin (0,04 mg/mL), neomycin (0,08 mg/mL), metronidazole (0,08 mg/mL), gentamycin (0,08 mg/mL) and sugar (15,0 mg/mL). Daily water consumption was roughly 5 mL/mouse.

### Antibody treatment

IgG1 α-MIF monoclonal antibodies were purified from ascitic fluid of mice injected with hybridomas (Clone NIHIII.D950) in the peritoneal cavity. Antibodies were purified using the NAb™ Protein A/G Spin Kit column (Thermo Scientific # 89980) and dialyzed using the Spectrum™ Spectra/Por™ Molecular Porous Membrane Tubing system (Thermo Scientific). To confirm antibody purification, samples were separated by SDS-PAGE, and stained with Coomassie Blue. The presence of only two bands referring to the light and heavy chains of the antibodies were observed. Samples were quantified using the commercial DCTM Protein Assay kit (Bio-Rad #500-0113 #500-0114) and filtered through 0.22 µm membranes. Mice were intraperitoneally treated daily with α-MIF IgG1 antibody (2,5 mg/kg) or IgG1 isotype control (2,5 mg/kg) starting one day before co-housing onset, and the treatment was maintained throughout the experiment.

### Intestinal cell isolation

For epithelium-lamina propria dissociation, feces were gently removed from the colon and the tube was cut transversely into three sections and then longitudinally. Specimens were rinsed with ice-cold PBS and incubated for 10 min with PBS-HEPES solution supplemented with DTT (1 mM) at 4 °C under agitation. Specimens were washed twice with PBS-HEPES solution supplemented with EDTA (30 mM) for 10 min at 37 °C under agitation. Tissue segments were transferred to a petri dish and the epithelium was mechanically separated from the lamina propria. Cells in the epithelial fraction were filtered through 100 µm cell strainers. The fraction corresponding to the lamina propria underwent enzymatic digestion in complete RPMI (cRPMI) (10% Fetal Bovine Serum (FBS), 1% PenStrep, 1% Sodium Pyruvate, 50µM β-mercaptoethanol, 1x HEPES) (RPMI-1640 with L-glutamine and sodium bicarbonate, Sigma #R8758) supplemented with Collagenase IV (1 mg/mL) (Worthington, #LS004188) and DNAse I (0.1 mg/mL) (Sigma, #SLCB2343) for 1h at 37 °C under shaking. Then, digested tissue was filtered through 100 µm cell strainers. Cell suspensions derived from both compartments were washed twice with PBS and subjected to 40% and 80% Percoll gradient enrichment (Percoll PLUS, GE Healthcare, #17-5445-01). Isolated cells were resuspended in cRPMI for counting. Mesenteric lymph nodes were readily macerated in cRPMI medium using 100 µm cell strainers. Cells were washed twice and kept in cRPMI medium, at 4 °C, until plating.

### Flow cytometry

Cells were transferred to 96-well U-bottom plates, centrifuged (400 g, 4 min, 4 °C) and the supernatant discarded. For *in vitro* restimulation, cells were incubated for 4 h at 37 °C with cRPMI supplemented with PMA (40 ng/mL) (Sigma #P8139), Ionomycin (1.5 µg/mL) (Sigma #IO634) and Brefeldin A (Biolegend #420601). Then, cells were washed and incubated with Live/Dead cell viability tracker (Invitrogen, #L34959) (1:1000) and the Fc-block CD16/CD32 Monoclonal Antibody (eBioscience, 14-0161-85) (1:1000) in PBS for 30 min at 4 °C. Cells were washed with PBS 2% FBS and incubated with surface antibody mix for 30 min in the dark at 4 °C. For intracellular staining, cells were fixed and permeabilized using eBioscience™ Foxp3 / Transcription Factor Staining Buffer Set (Invitrogen, #00-5523-00). Cells were incubated with an intracellular antibody mix for 30 min in the dark at 4 °C. The following antibodies were used: α-CD4 PERCP (BD #553052, clone RM4-5), α-CD8α e780 (eBioscience #47-0081-82, clone 53-6.7), α-Foxp3 e450 (eBioscience #48577380, clone FJK-165), α-IFNγ FITC (BD #554411, clone E7), α-IL-17A AF647 (BD, #560184, clone TC11-18H10), α-TNFα PE (eBioscience #12-7321-81, clone MP6-XT22), α-CD11b APCe780 (Invitrogen #47-0112-82, clone M1/70), α-Ly6G BV605 (BD #563005, clone 1A8), α-SiglecF AF647 (BD #562680, clone E50-2440), α-Ly6C PERCP Cy5.5 (BD #128011, clone HK1.4), α-F4/80 PE, α-CD11c FITC (BD #553801, clone HL3). Samples were acquired on the BD LSR Fortessa™ cytometer and analyzed using FlowJo v10.0.7 software.

### Flow cytometry of fecal bacteria

For bacterial IgA/IgM coating experiments, 8 mg of fresh feces were collected, macerated in 1 mL of PBS and centrifuged at low speed (500 g, 5 min, 4 °C). The supernatant was transferred to a new tube, washed with PBS 1% BSA and centrifuged at high speed (12000 g, 5 min, 4 °C). The pellet was resuspended in blocking solution (PBS 1% BSA 20% rat serum) for 20 min at 4 °C. Cells were washed twice and resuspended in antibody mix solution containing α-IgM PE (Jackson Immuno Research #115-116-075) and α-IgA BV605 (BD OptiBuild #743295, clone C10-1) for 30 min in the dark at 4 °C. Samples were washed twice and fixed with fresh paraformaldehyde (4%) for 30 min at 4°C. Bacteria were washed, kept in PBS at 4 °C and protected from light until acquisition. Bacteria were resuspended in PBS-DAPI solution immediately before acquisition in the cytometer.

### Bacteria cultivation

Fresh fecal samples were collected from AVNM-treated *Il10^-/-^*mice after seven days of co-housing with AVNM-treated *Mif^-/-^*mice. Bacteria were cultivated anaerobically in Brain Heart Infusion (BHI) agar supplemented with 10% horse blood. For gentamycin sensibility assay *in vitro,* BHI agar was supplemented or not with gentamycin (200 μg/ mL). *K. oxytoca* and *P. mirabilis* were aerobically cultured in BHI agar for 18h. Bacteria suspensions were prepared in PBS (OD_600_ = 0,6: *K. oxytoca* ≅ 1 x 10^10^ CFU/ mL and *P. mirabilis* ≅ 1 x 10^10^ CFU/ mL). Mice were mono-inoculated with 1 x 10^9^ CFU or co-inoculated with 0.5 x 10^9^ CFU of each species through oral gavage.

### Stool DNA extraction and 16S rRNA sequencing

Fresh stool samples were collected from WT, *Mif^-/-^* and *Il10^-/-^* mice at steady state and stored at -80°C. Fecal DNA extraction was performed using the QIAamp PowerFecal Pro DNA kit (Qiagen, Dusseldorf, Germany) according to the previously described method. The total bacterial DNA was eluted with 100μl of elution buffer and stored at -20°C before PCR amplification. The concentration and purity of the isolated DNA samples were analysed in NanoDrop® (Thermo Scientific). The isolated DNA was subjected to PCR amplification, targeting the V5 and V6 hypervariable regions of the 16S rRNA gene. We employed bacteria-specific primers, with sequences previously described: forward 5’-CCATCTCATCCCTGCGTGTCTCCGACTCAGC barcode ATTAGATACCCYGGTAG TCC-3’ and reverse 5’-CCTCTCTATGGGCAGTCGGTGATACGAGCTGACGACARC CATG-3’.The thermal cycling conditions were set as follows: an initial denaturation at 94°C for 5 minutes, followed by 35 cycles consisting of denaturation at 94°C for 1 minute, annealing at 46°C for 20 seconds, and elongation at 72°C for 30 seconds, and a final elongation phase at 72°C for 7 minutes. This protocol ensures the specific amplification of the bacterial 16S rDNA regions of interest (Yilmaz et al., 2019). The PCR-generated amplicons were resolved using a 1% agarose gel, displaying an anticipated amplicon size of approximately 350 base pairs. Following electrophoresis, the amplicons were purified utilizing the QIAQuick Gel Extraction Kit (Qiagen, Dusseldorf, Germany). Quantification of the purified amplicons was performed using the Qubit dsDNA HS Assay Kit on the Qubit 3.0 Fluorometer (ThermoFisher Scientific). For the preparation of template-positive Ion PGM™ Template OT2 400 Ion Sphere™ Particles (ISPs) harboring clonally amplified DNA, we employed the Ion OneTouch™ Instrument alongside the Ion PGM™ Template OT2 400 Kit (ThermoFisher). Subsequently, sequencing was executed using the Ion PGM™ Sequencing 400 Kit and Ion 316™ Chip V2, within the framework of the Ion PGM™ System (Thermo Fisher) (Whiteley et al., 2012).

### Stool microbiota analysis

The fastq sequencing files produced by the Ion Torrent PGM™ System were imported into the Quantitative Insights into Microbial Ecology (QIIME), consistent with methodologies as described before (Caporaso et al., 2010). Data analysis was performed using the QIIME pipeline. Operational taxonomic units were picked using UCLUST with a 97% sequence identity threshold followed by taxonomy assignment using the latest Greengenes database (http://greengenes.secondgenome.com/downloads). The calculation of species richness (alpha diversity metrics), distance between samples/groups in terms of microbial composition (beta diversity) and taxonomy analysis were performed using the *phyloseq* package in R, version 3.0.2 (http://www.R-project.org) (Callahan et al., 2016; McMurdie and Holmes, 2013). The non-parametric Mann-Whitney U-tests for alpha diversity and Adonis (Permanova method on distances; 9999 permutations) for beta diversity analysis were used to assess statistical differences between groups with p<0.05 considered significant. Multivariate analysis by linear models (MaAsLin2) R package was used to analyze the taxa present in at least 30% of the samples and OTUs that had more than %0.001 of total counts (Morgan et al., 2012). Data with q<0.05 (adj-p value; Benjamini-Hochberg false discovery rate correction) were considered significant.

### RNA extraction and gene expression

For colon RNA extraction, approximately 100 mg of clean, feces-free tissue was collected from WT and *Mif^-/-^* mice under homeostatic conditions and immediately macerated in the commercial Fastzol reagent (Quatro G Biotechnology, #100044). Briefly, after maceration, the samples were incubated with chloroform for 3 min, centrifuged (15 min, 12000g, 4 °C), and the upper clear aqueous phase was collected into a new tube. The enriched RNA fraction was incubated with isopropanol for 10 min at 4 °C, centrifuged (10 min, 12000g, 4 °C), and the supernatant was discarded. The RNA precipitate was resuspended in 75% ethanol, centrifuged (5 min, 7500g, 4°C) and the supernatant discarded. The tubes containing the samples were left open to evaporate residual ethanol, and the samples were resuspended in nuclease-free water. RNA was quantified using the NanoDrop 2000 reader (Thermo Scientific) by measuring absorbances at 260 nm, and purity was determined based on 260/280 and 260/230 ratios. Reverse transcription (RT) was performed using the commercial High Capacity RNA-to-cDNA kit (Applied Biosystems, #4388950) on samples previously adjusted to a concentration of 0.1 μg/μL. Quantitative polymerase chain reaction (qPCR) was prepared in 96-well plates using the commercial mix SYBR™ Green PCR Master Mix (Applied Biosystems, #4309155) and the following specific primers: *Gapdh* FW 5’-AGGTCGGTGTGAACGGATTTG-3’, RV 5’-TGTAGACCATGTAGTTGAGGTCA-3’; *Tnfa* FW 5’-CCTCACACTCAGATCATCTTCTCA-3’, RV 5’-TGGTTGTCTTTGAGATCCATGC-3’; *Il6* FW 5’-TCATATCT TCAACCAAGAGGTA-3’, RV 5’-CAGTGAGGAATGTCCACAAACTG-3’. Reaction occurred in the 7500 Real-Time PCR System thermocycler (Applied Biosystems). Relative quantification (RQ) was performed by the 2^−ΔΔCt^ method using *Gapdh* as the endogenous gene.

### Statistical analysis

Mice body weight comparisons were performed by calculating the area under the curve (AUC). Unless otherwise stated in the text, comparisons between two groups were made with a two-tailed T-test and comparisons between more than two groups with one-way ANOVA followed by post-hoc Tukey’s test. Survival curves analyses were performed using log-rank Mantel-Cox test. * for adjusted p ≤ 0.05; ** for adjusted p ≤ 0.01; *** for p ≤ 0.001; **** for p ≤ 0.0001. Statistical analyses were performed using the software GraphPad Prism version 10.0.0 for Windows (GraphPad Software, Boston, Massachusetts, USA).

## Acknowledgments

We would like to thank Prof. Dr. João C. Machado (COPPE/UFRJ) and Dr. Rodrigo P. Oliveira (COPPE/UFRJ) for the support during colonoscopy experiments. We thank Prof. Dr. Renata C. Picão (IMPG/UFRJ) and Dr. Luana Boff (IMPG/UFRJ) for kindly assisting with *K. oxytoca* and *P. mirabilis* cultivation. We thank Dr. Letícia S. Alves for animal care and experimental support. We thank the microscopy platform Unidade de Microscopia Multiusuário Padrón-Lins (UNIMICRO) (IMPG/UFRJ) and Dr. Jefferson B. S. Cypriano for the assistance. We also thank veterinarian MSc. Isabel Roussouliéres (CCS/UFRJ) and Instituto de Ciência e Tecnologia em Biomodelos (ICTB/Fiocruz) for the mice health surveillance in our animal facility. The hybridoma for the production of α-MIF antibody was generously provided by Dr. Richard Bucala (Yale University). We thank SciDraw for scientific drawings. This work was supported by grants from Coordenação de Aperfeiçoamento de Pessoal de Nível Superior (CAPES), Conselho Nacional de Desenvolvimento Científico e Tecnológico (CNPq), by Fundação de Amparo à Pesquisa do Estado do Rio de Janeiro (FAPERJ) and by a Fullbright fellowship to MTB. VMV, EMC and AMSG were supported by fellowships from CAPES. LMF was supported by fellowships from CAPES and FAPERJ.

## Author’s contribution

Conducted experiments: VMV, LMF, BY, EMC, AMSG, JNDF, VR, JZK, SGR, GAT, HSPS, FBC

Supervised and designed experiments: VMV, FBC, MTB

Data analysis: VMV, BY, EMC, VR, HSPS

Experimental result interpretation: VMV, BY, EMC, DL, HSPS, FBC, MTB

Manuscript writing: VMV, FBC, MTB

Funding acquisition: MTB

## Data availability statement

The data that support the findings of this study are available on request from the corresponding author.

---

**Supplementary Table 1.**
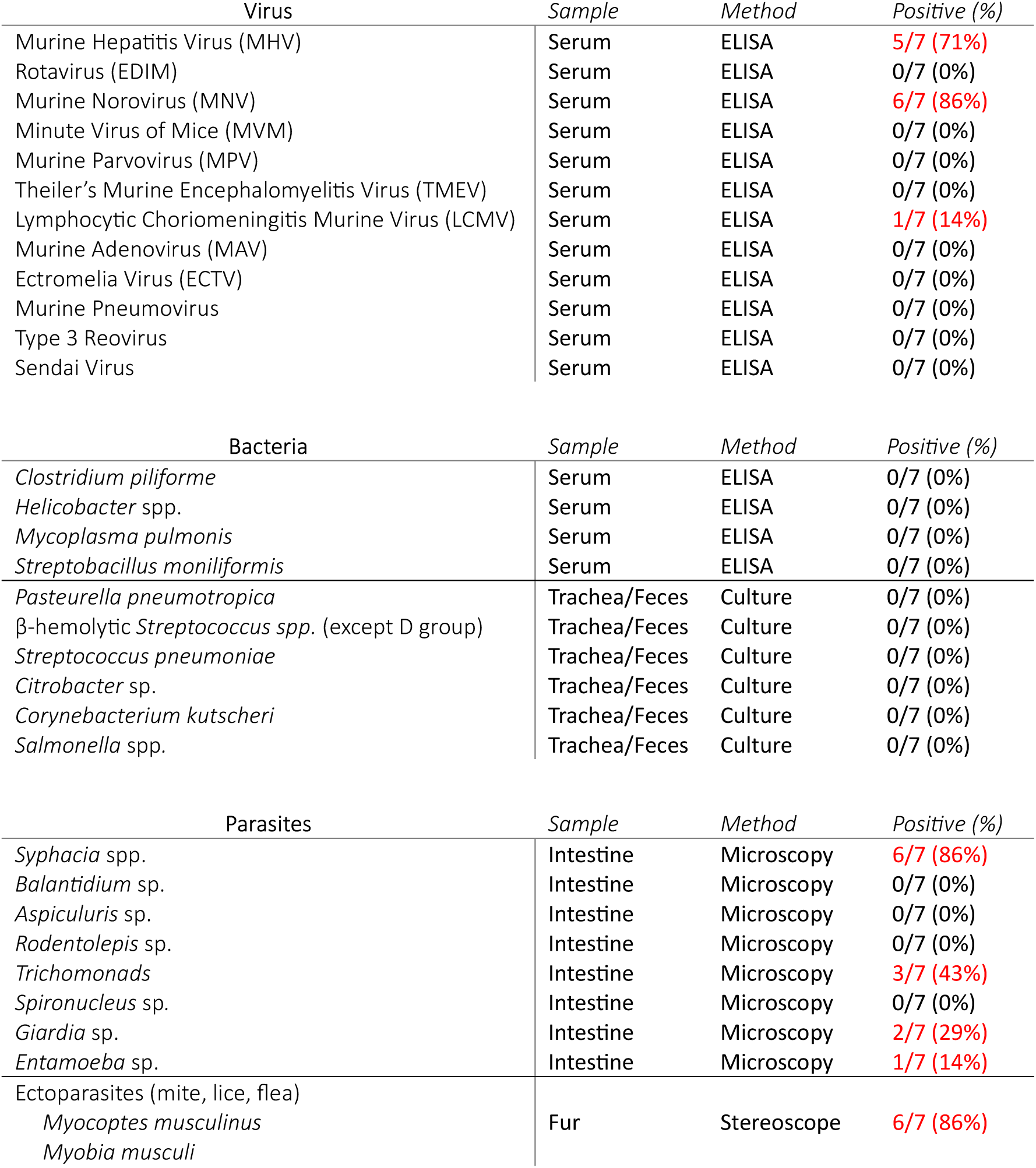
Health monitoring of mice kept in the vivarium of the Inflammation and Immunity Laboratory (IMPG/UFRJ). WT (n=2), *Mif^-/-^* (n=4) and *Il10^-/-^* (n=1) mice at least six months old were selected from our vivarium and screened for a list of pathobionts, pathogens and parasites. Surveillance was conducted in 2022 and 2023.

**Extended Figure 1.**
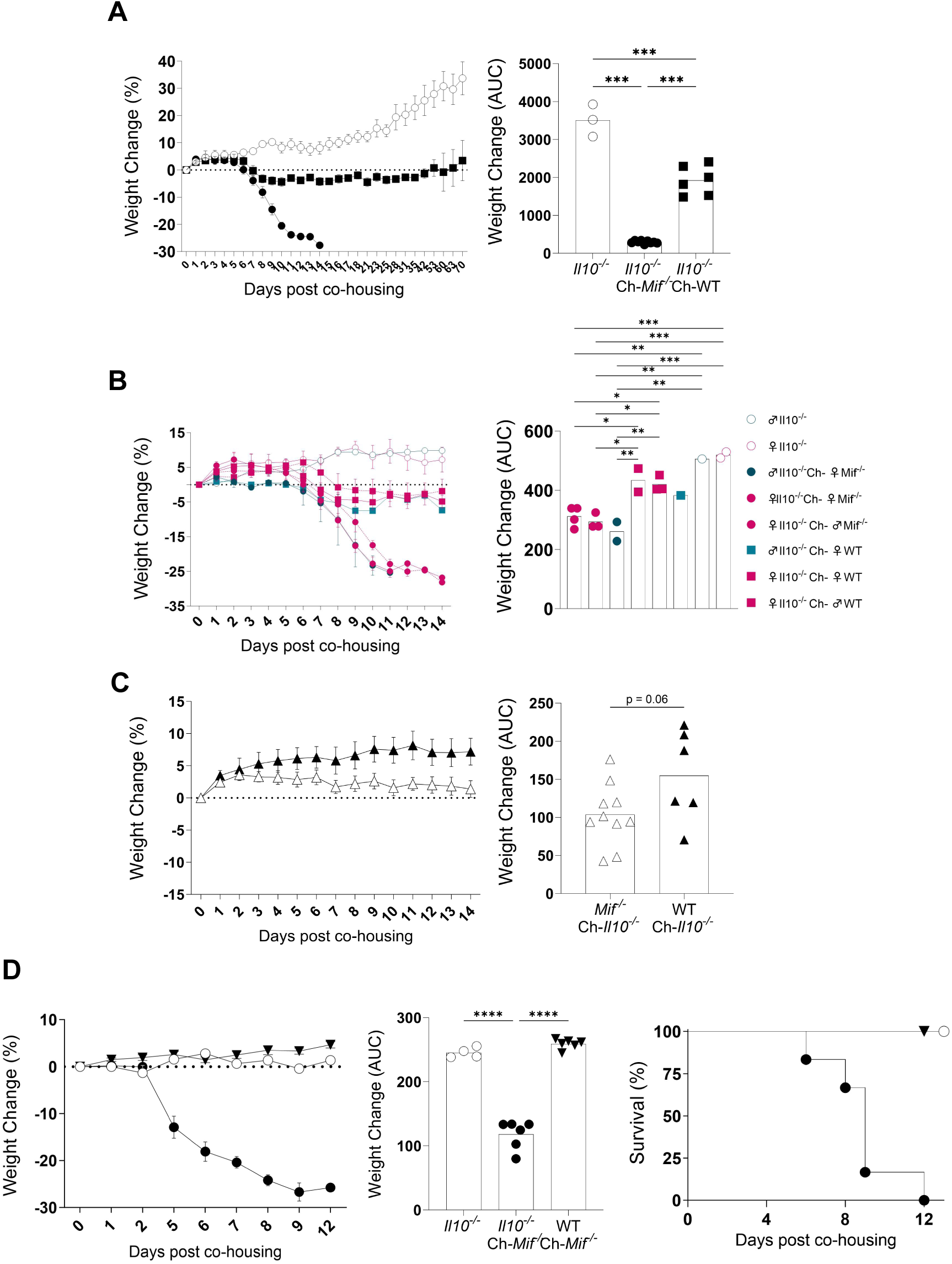
Different co-housing conditions reinforce that specifically *Il10^-/-^* mice develop lethal disease, and only when co-housed with *Mif^-/-^* mice. **(A-B)** The same experiment as in Figure 1B, **(A)** showing data from later timepoints – mice were followed up to 70 days after co-housing – and **(B)** separated by sex. **(C)** *Mif^-/-^* and WT mice body weight curve after co-housing *Il10^-/-^* mice. Two-tailed T-test. **(D)** *Il10^-/-^* and WT mice body weight (left) and survival (right) curves after co-housing *Mif^-/-^* mice. **(A,B,D)** One-way ANOVA, post-hoc Tukey. **(A-D)** Weight mean ± SEM. Survival curves were compared with log-rank Mantel-Cox test. * for adjusted p ≤ 0.05; ** for adjusted p ≤ 0.01; *** for p ≤ 0.001.

**Extended Figure 2.**
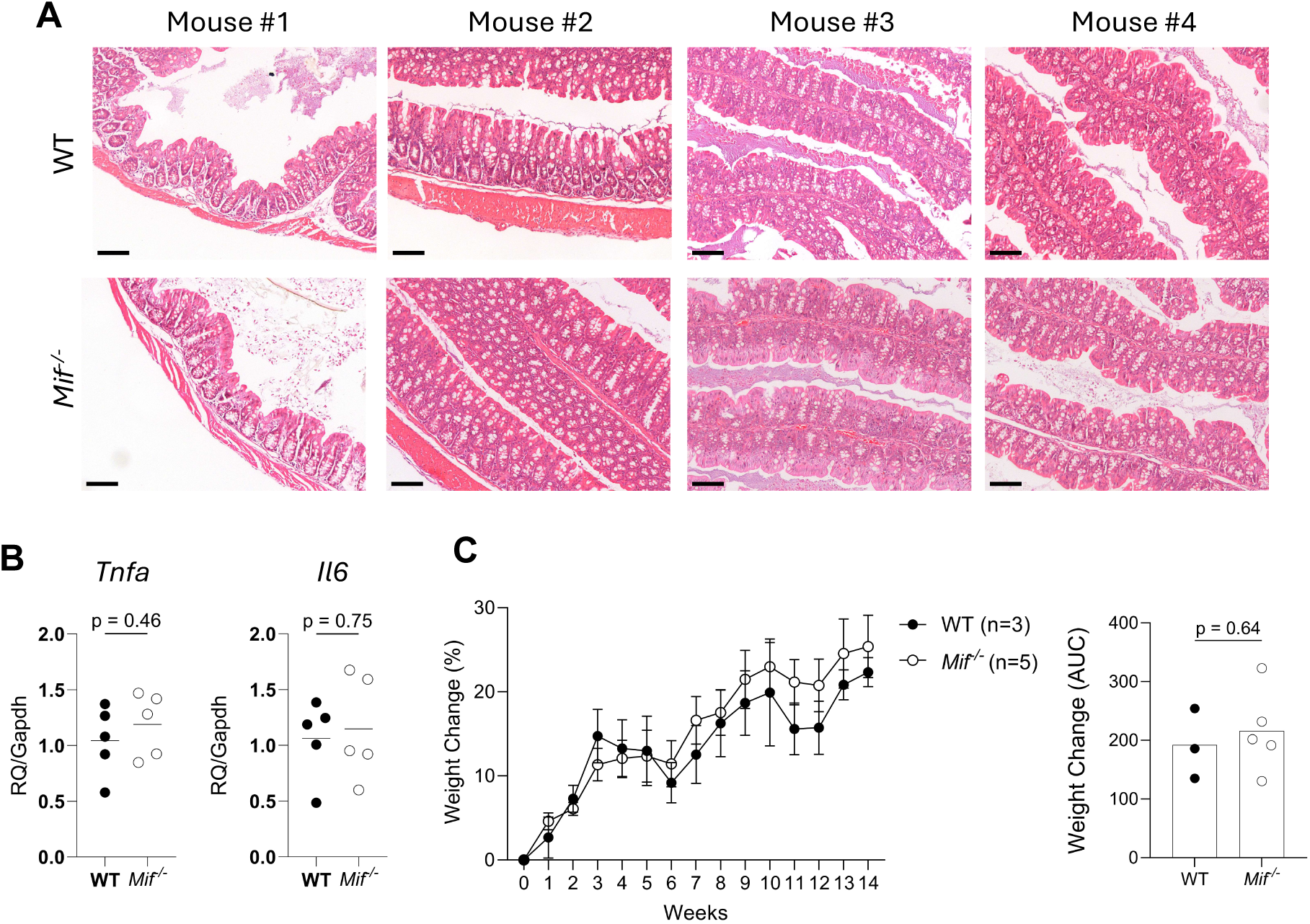
*Mif^-/-^* mice do not exhibit overt signs of intestinal inflammation during homeostasis. Colonic tissue samples were collected from *Mif^-/-^* and WT mice at steady state. **(A)** Representative colon photomicrographs from four mice in each group. Hematoxylin/Eosin (HE) staining was performed on 5 µm colon sections. Scale Bar = 100 µm; magnification = x100. **(B)** *Tnfa* and *Il6* gene expression assessed by RT-qPCR. Relative quantification (RQ) was performed by the 2^−ΔΔCt^ method using *Gapdh* as the endogenous gene. **(C)** Six-week-old *Mif^-/-^* and WT mice body weight curve over 14 weeks under a conventional diet. Weight mean ± SEM. **(B,C)** Two-tailed T-test.

**Extended Figure 3.**
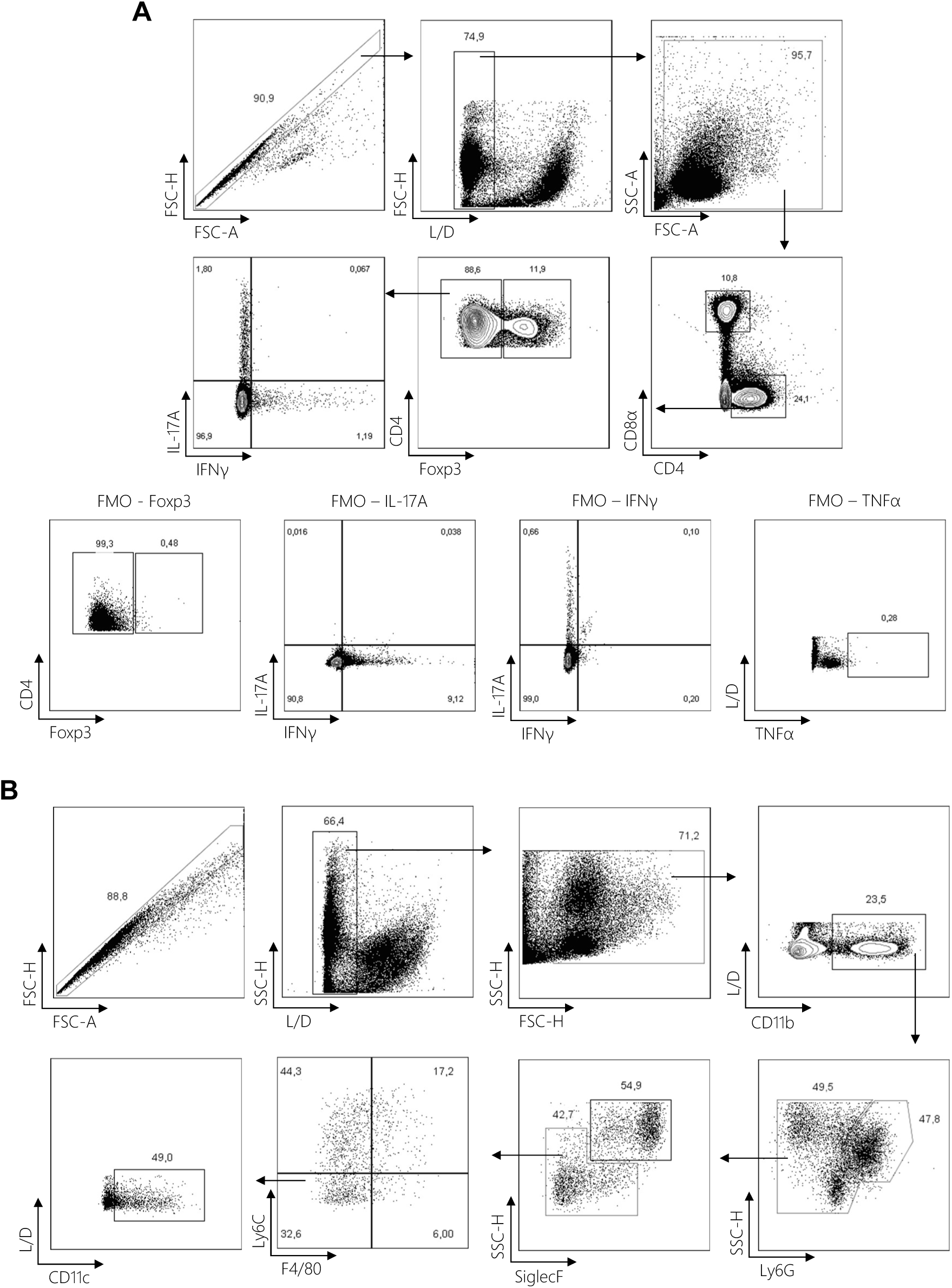
Flow cytometry gating strategy. Immunophenotyping of colonic lamina propria, colonic epithelium, and mesenteric lymph nodes was performed by flow cytometry. **(A)** lymphoid and **(B)** myeloid cells were identified after doublet, debris and dead cell exclusion. FMO, Fluorescence Minus One.

**Extended Figure 4.**
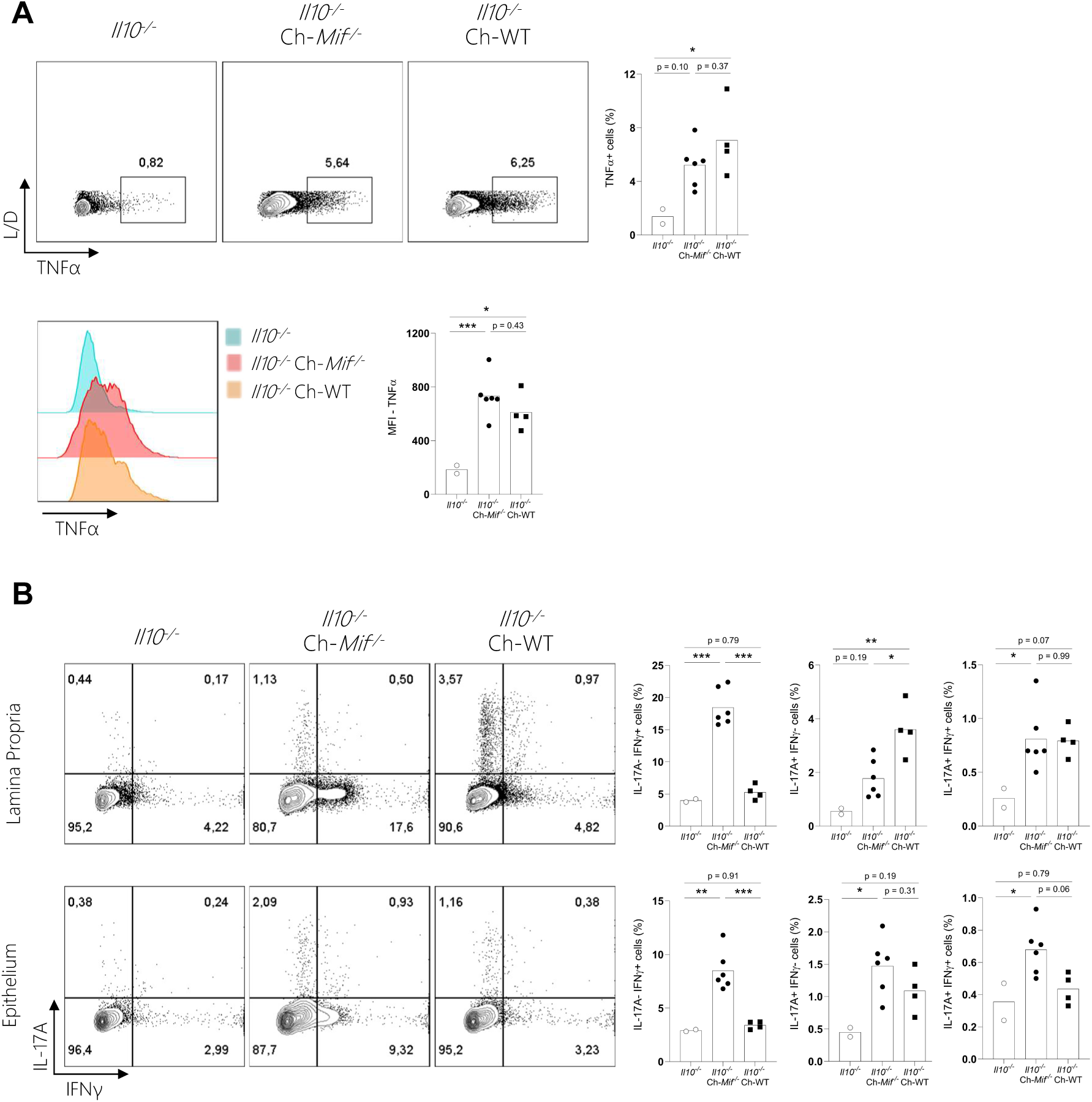
Increased IFNγ^+^ cells are exclusively observed in *Il10^-/-^* mice after co-housing with *Mif^-/-^* mice. **(A)** Immunophenotyping of *Il10^-/-^* mice colonic lamina propria after co-housing. Top: representative contour plots and column bar graph showing the frequency of TNFα^+^ cells. Bottom: representative histogram and column bar graph of TNFα Median Fluorescence Intensity (MFI). **(B)** Frequencies of IFNγ^+^ and IL-17A^+^ cells in *Il10^-/-^* mice colonic lamina propria and epithelium after co-housing. Representative contour plots. **(A and B)** Frequencies after doublet, debris and dead cell exclusion. One-way ANOVA, post-hoc Tukey. * for adjusted p ≤ 0.05; ** for adjusted p ≤ 0.01; *** for p ≤ 0.001.

**Extended Figure 5.**
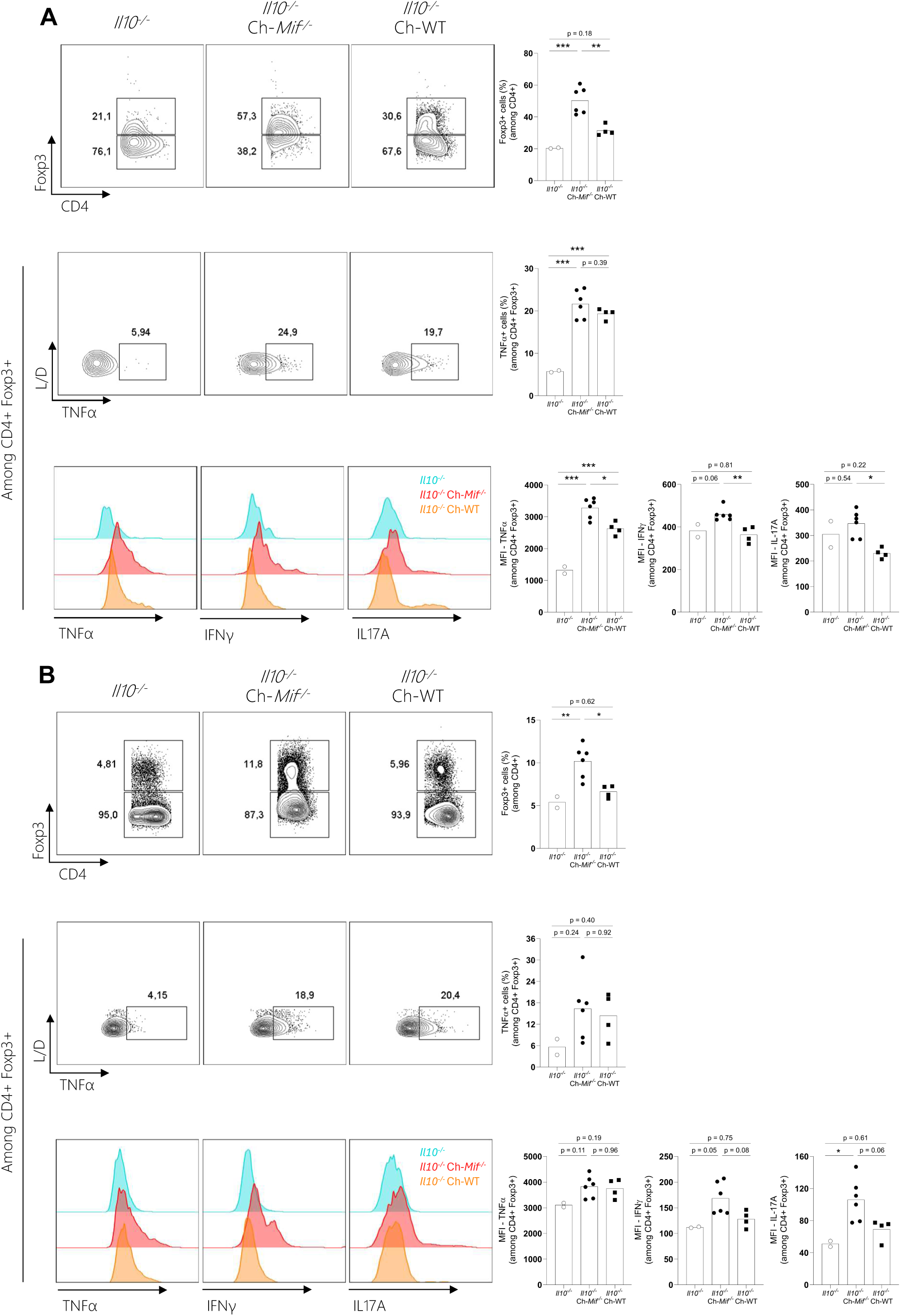
*Il10^-/-^* mice display increased, but phenotypically subverted, CD4^+^Foxp3^+^ Treg cells after co-housing with *Mif^-/-^* mice. Immunophenotyping of *Il10^-/-^* mice **(A)** colonic lamina propria and **(B)** mesenteric lymph nodes after co-housing. Representative contour plots and column bar graphs showing the frequencies of CD4^+^Foxp3^+^ Treg cells. Representative contour plots and column bar graphs showing the frequencies of TNFα^+^ cells among CD4^+^Foxp3^+^ Treg cells. Representative histogram and column bar graph of TNFα, IFNγ and IL-17A Median Fluorescence Intensity (MFI) among CD4^+^Foxp3^+^ Treg cells. One-way ANOVA, post-hoc Tukey. * for adjusted p ≤ 0.05; ** for adjusted p ≤ 0.01; *** for p ≤ 0.001. MFI, Median Fluorescence Intensity.

**Extended Figure 6.**
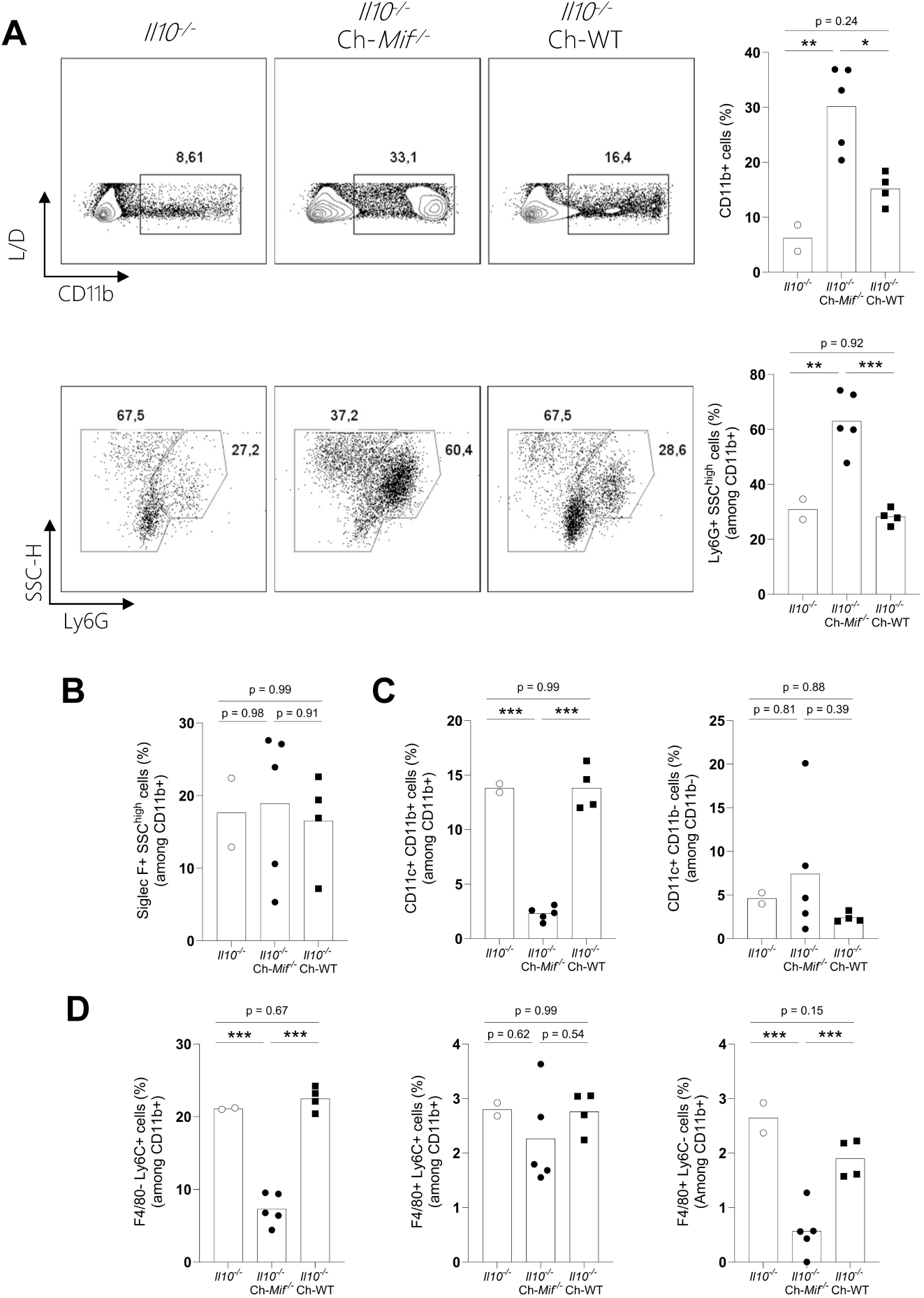
*Il10^-/-^* mice exhibit neutrophilic infiltrate in the colon after co-housing with *Mif^-/-^* mice. Relative frequencies of different myeloid populations in *Il10^-/-^* mice colonic lamina propria after co-housing. **(A)** Top: frequency of CD11b^+^ myeloid cells relative to live cells (after doublet, debris and dead cell exclusion). Bottom: frequency of CD11b^+^Ly6G^+^SSC^high^ neutrophils relative to CD11b^+^ cells. **(B)** Frequency of CD11b^+^Ly6G^-^SiglecF^+^SSC^high^ eosinophils, **(C)** CD11b^+^Ly6G^-^SiglecF^-^Ly6C^-^F4/80^-^CD11c^+^ dendritic cells, and **(D)** CD11b^+^Ly6G^-^SiglecF^-^Ly6C^+^F4/80^-^ monocytes and CD11b^+^Ly6G^-^SiglecF^-^F4/80^+^ F4/80+ macrophages relative to CD11b^+^ cells. For the gating strategy, see Figure S2. One-way ANOVA, post-hoc Tukey. * for adjusted p ≤ 0.05; ** for adjusted p ≤ 0.01; *** for p ≤ 0.001.

**Extended Figure 7.**
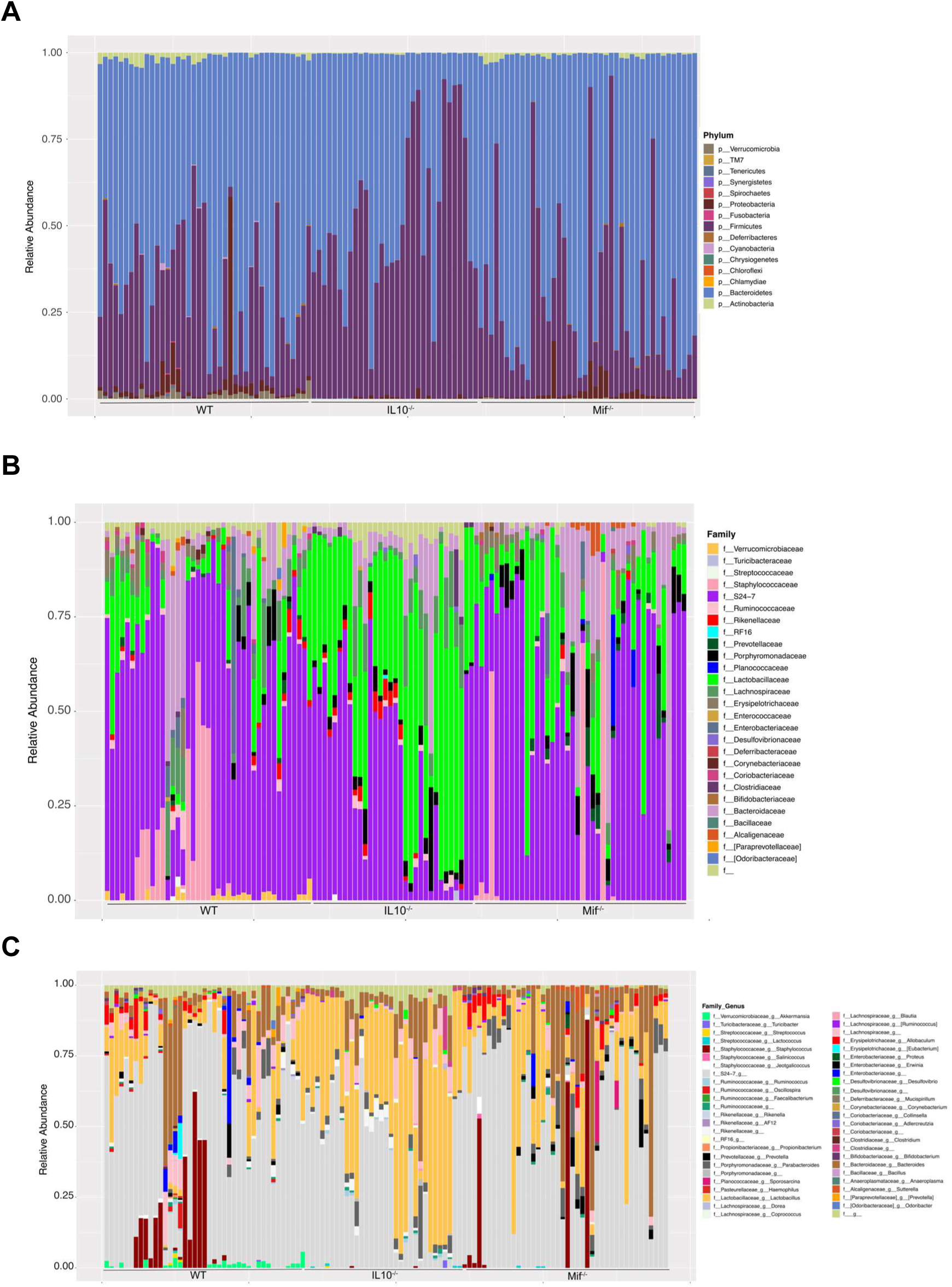
*Mif^-/-^* mice harbor a unique intestinal microbiota. Stool samples from *Mif^-/-^*, *Il10^-/-^* and WT mice under steady state conditions were analyzed for microbiome composition based on the 16S rRNA gene sequencing. Relative abundance at **(A)** phylum, **(B)** family and **(C)** genus level.

**Extended Figure 8.**
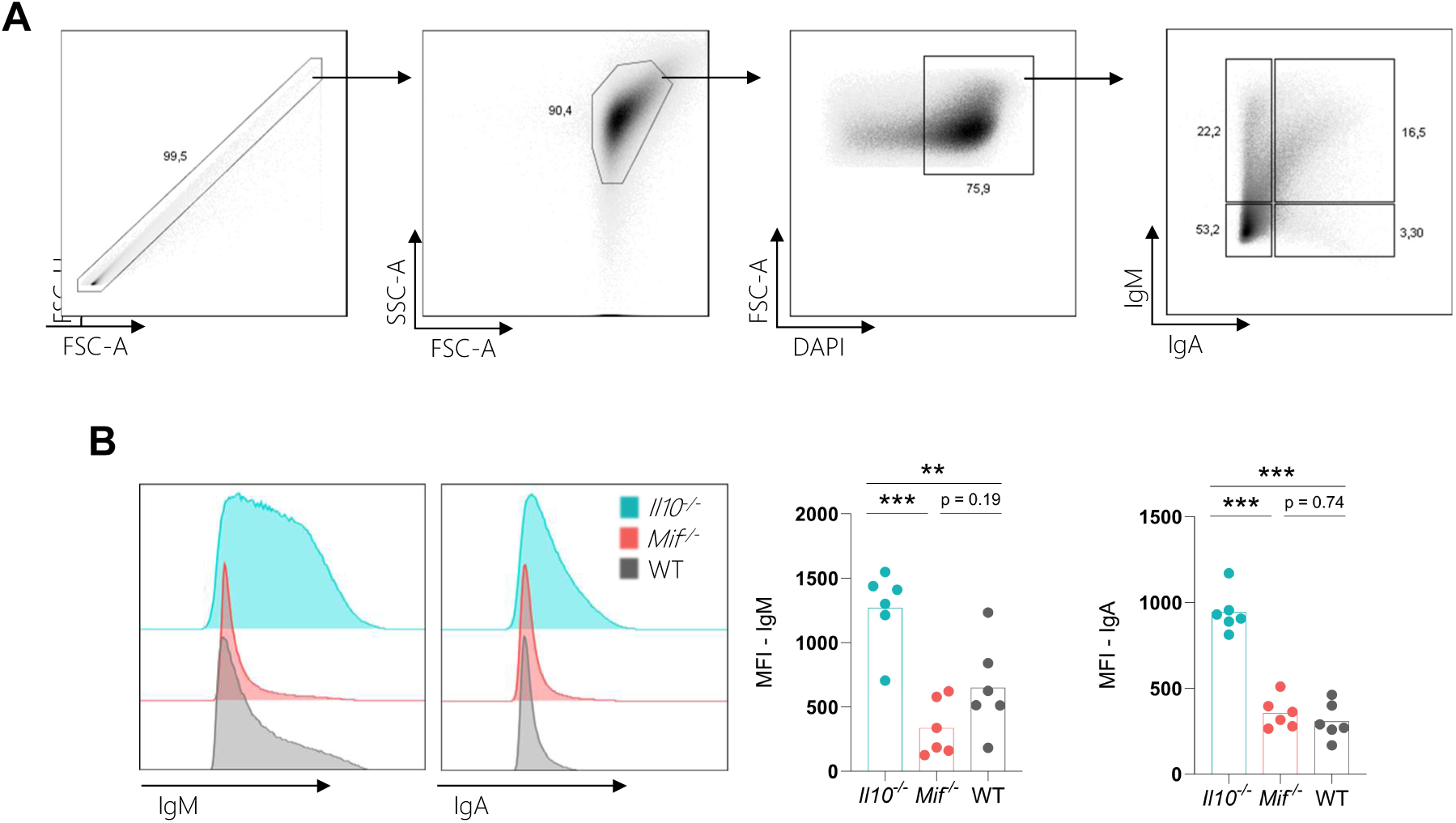
*Mif^-/-^* mice shelter an increased community of antibody uncoated bacteria. Bacterial cells were isolated from fresh fecal pellets from *Mif^-/-^*, *Il10^-/-^* and WT mice under steady state conditions. **(A)** The gating strategy depicts doublet exclusion and bacterial cells selected by size and DAPI staining. **(B)** Representative histogram and column bar graphs of IgM and IgA Median Fluorescence Intensity (MFI). One-way ANOVA, post-hoc Tukey. ** for adjusted p ≤ 0.01; *** for p ≤ 0.001.

**Extended Figure 9.**
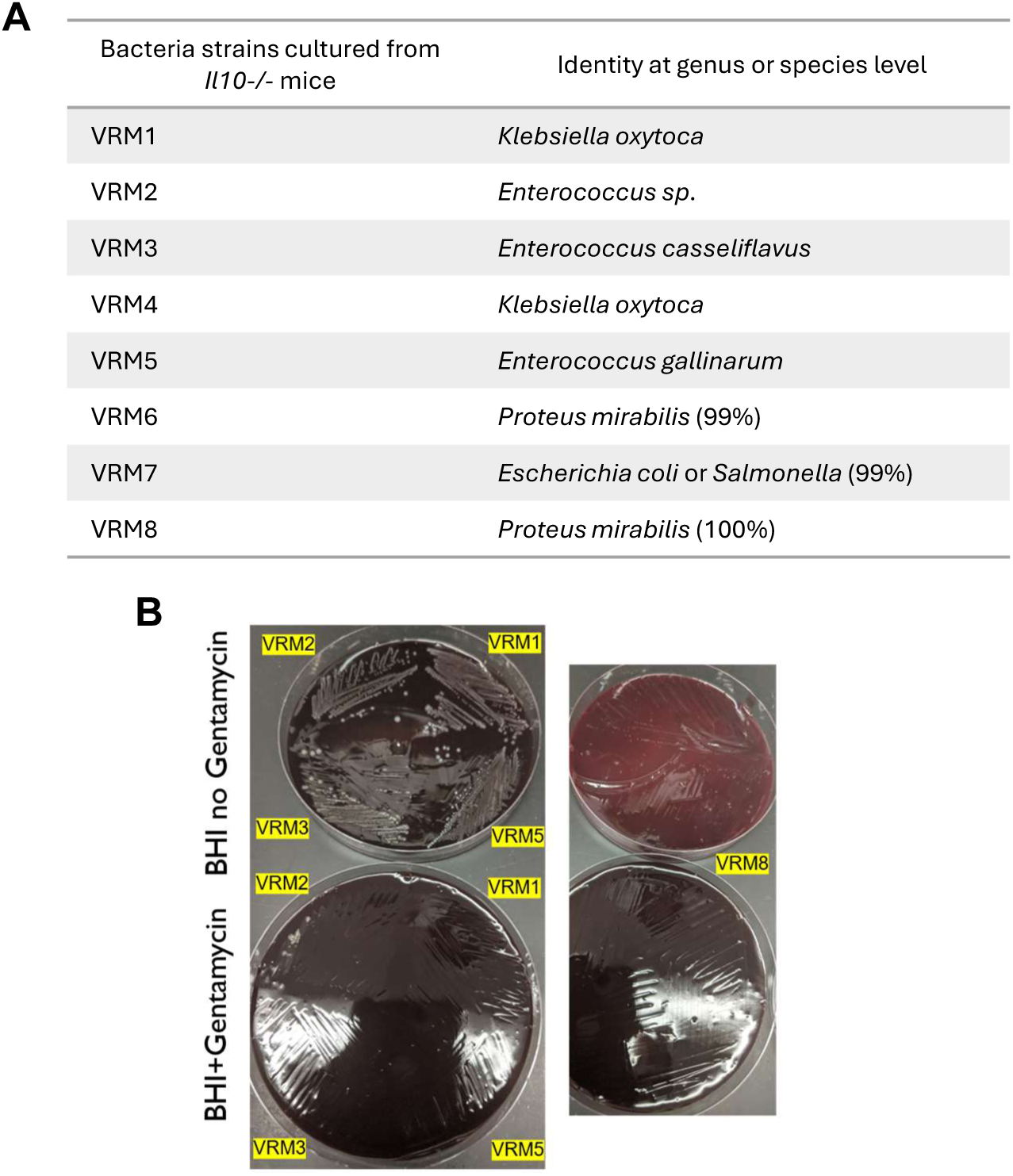
*Klebsiella oxytoca, Proteus mirabilis* and *Enterococcus* spp. are putative candidates for triggering colitis in *Il10^-/-^* mice. Fresh fecal pellets were collected from *Il10^-/-^* mice under the AVNM regime and after co-housing *Mif^-/-^* mice for seven days. **(A)** Fecal bacteria were cultivated under anaerobe conditions in BHI agar supplemented with 10% horse blood. Individual colonies were isolated by morphology and identified based on partial 16s rRNA gene sequencing. **(B)** Isolated strains cultivated in BHI media supplemented or not with gentamycin.

**Extended Figure 10.**
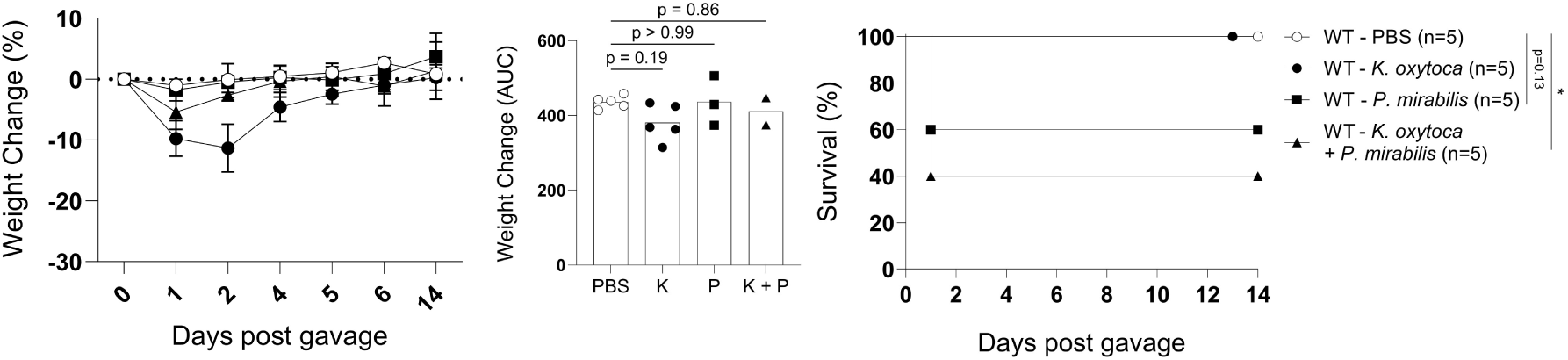
*K. oxytoca* and *P.mirabilis* are less lethal to WT mice. WT mice were mono-inoculated through oral gavage with 1 x 10^9^ CFU of *P. mirabilis*, 1 x 10^9^ CFU of *K. oxytoca,* or co-inoculated with 0.5 x 10^9^ CFU of each species. Mice body weight (left) and survival (right) curves after inoculation. Weight mean ± SEM. One-way ANOVA, post-hoc Tukey test for weight change (AUC) and log-rank Mantel-Cox for survival curve.

**Extended Figure 11.**
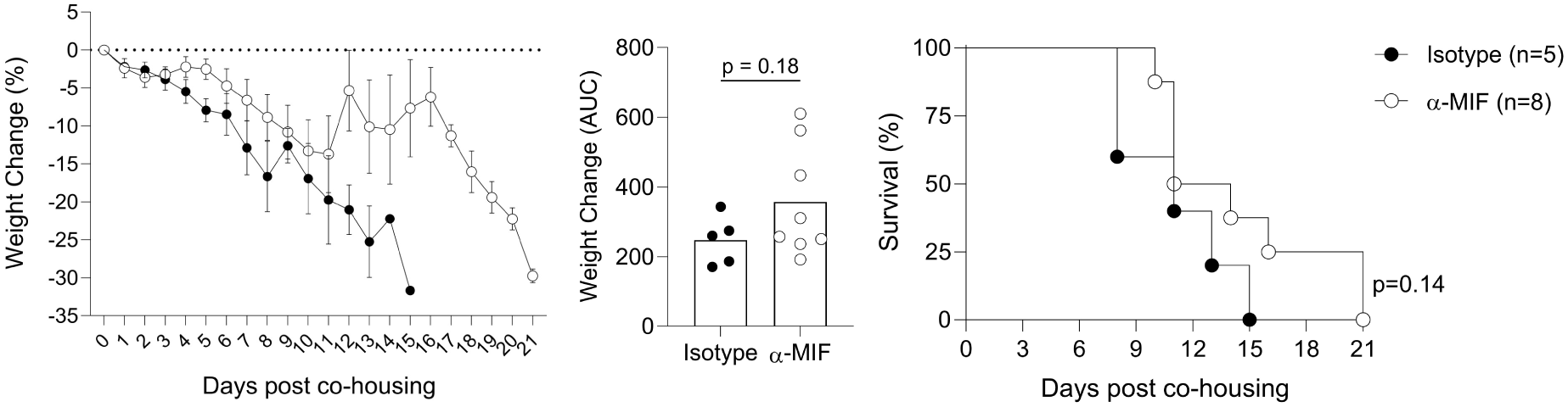
MIF blockade fails to rescue *Il10^-/-^* mice from lethal co-housing with *Mif^-/-^* mice. *Il10^-/-^* mice body weight (left) and survival (right) curves after co-housing with *Mif^-/-^* mice. *Il10^-/-^* mice were treated intraperitoneally with 2,5 mg/kg of α-MIF monoclonal antibody or IgG1 isotype control daily. Treatment started the day before co-housing onset, and was maintained throughout the experiment. Mean ± SEM. Pool of two independent experiments. Two-tailed T-test for weight change (AUC) and log-rank Mantel-Cox for survival curve.

